# A structural and kinetic link between membrane association and amyloid fibril formation of α-Synuclein

**DOI:** 10.1101/173724

**Authors:** Thibault Viennet, Michael M. Wördehoff, Boran Uluca, Chetan Poojari, Hamed Shaykhalishahi, Dieter Willbold, Birgit Strodel, Henrike Heise, Alexander K. Buell, Wolfgang Hoyer, Manuel Etzkorn

## Abstract

The protein α-Synuclein (αS) is linked to Parkinson’s disease through its abnormal aggregation, which is thought to involve an interplay between cytosolic and membrane-bound forms of αS. Therefore, better insights into the molecular determinants of membrane association and their implications for protein aggregation may help deciphering the pathogenesis of Parkinson’s disease. Following previous studies using micelles and vesicles, we present a comprehensive study of αS interaction with phospholipid bilayer nanodiscs. Using a combination of NMR - spectroscopic and complementary biophysical as well as computational methods we structurally and kinetically characterize αS interaction with defined stable planar membranes in a quantitative and site-resolved way. We probe the role of αS acetylation as well as membrane charge, plasticity and available surface area in modulating αS membrane binding modes and directly link these findings to their consequences for αS amyloid fibril formation.

---

The protein α-Synuclein (αS) is associated with various synucleinopathies including Parkinson’s disease through its abnormal aggregation, fibril formation and formation of Lewy bodies (Jucker and Walker, 2013, Luk, et al., 2012, Spillantini, et al., 1997, Tuttle, et al., 2016). While the exact native function is yet not fully understood, αS is found in synaptic vesicles and supposed to be involved in membrane interactions, e.g. in synaptic vesicle homeostasis (Bellani, et al., 2010, Gitler, et al., 2008) and SNARE-like vesicle-to-vesicle or vesicle-to-membrane fusion (Diao, et al., 2013, Fusco, et al., 2016). Membrane association of αS has been shown to modulate its aggregation propensity (Dikiy and Eliezer, 2012, Zhu, et al., 2003), and αS oligomeric species have been proposed to be the toxic species in Parkinson’s disease, especially through membrane pore formation mechanisms (Auluck, et al., 2010, Butterfield and Lashuel, 2010). αS has been shown to be N-terminally acetylated, which is thought to act as an important mode of regulation of protein-membrane association (Dikiy and Eliezer, 2014, Nemani, et al., 2010, Theillet, et al., 2016).

Previous data recorded using micelle and vesicle preparations already provided valuable information of the αS-membrane interactions, including binding and lipid specificity (Bodner, et al., 2009, Jo, et al., 2000, Rhoades, et al., 2006), effect of mutations on membrane association (Bodner, et al., 2010, Fusco, et al., 2016), micelle-bound structure (Ulmer, et al., 2005), vesicle-bound structural insights (Drescher, et al., 2008, Fusco, et al., 2014, Jao, et al., 2008) and conformational dynamics (Eliezer, et al., 2001, Fusco, et al., 2014, Fusco, et al., 2016). Two structural models of lipid-bound αS were proposed, i.e. the “extended helix” consisting of one roughly 100-residue long α-helix (Georgieva, et al., 2008) and the “horse-shoe”, consisting of two helices with different lipid affinities separated by a kink at residues 42-44 (Drescher, et al., 2008). Furthermore, various effects of lipids for αS aggregation were reported including inhibition of aggregation (Zhu and Fink, 2003), triggering of fibrillation (Galvagnion, et al., 2015, Zhao, et al., 2004) and modification of fibril structure (Comellas, et al., 2012). Membrane binding and its effect on aggregation have been shown to be strongly dependent on chemical properties of the lipids including head group charge content (Drescher, et al., 2008) and fatty acid type (Galvagnion, et al., 2016).

The intrinsic features of the phospholipid bilayer nanodisc (ND) system (Bayburt, et al., 2002) offer the potential to provide additional insights that are complimentary to the information obtained on micelle and vesicle preparations. Notably, NDs have been used before to study the effect of calcium ions on the membrane interaction of αS (Zhang, et al., 2014) as well as lipid and monomer specificity of the Alzheimer’s associated Aβ peptide (Thomaier, et al., 2016). In general, NDs are very homogenous, stable in a wide buffer range (Viegas, et al., 2016) and allow the preparations of well-defined lipid mixtures with an accurate estimate of the bilayer size (Denisov, et al., 2004), charge (Her, et al., 2016), and lipid molarity (Inagaki, et al., 2012). The increased stability offers e.g. the possibility to consistently determine the interaction with a stable planar bilayer surface. In contrast, it is known for small unilamellar vesicles (SUVs) that the interaction with αS can considerably and rapidly change the lipid environment (e.g. from homogenous SUVs to rather heterogeneous particles (Comellas, et al., 2012, Eichmann, et al., 2016, Fusco, et al., 2016, Ouberai, et al., 2013)). Additionally, the smaller size of the NDs should, in theory, allow the detection of the lipid-bound state using suitable solution NMR techniques (Etzkorn, et al., 2013, Hagn, et al., 2013, Viegas, et al., 2016). In general, the well-defined size and lipid composition of NDs, paired with their accessibility, homogeneity and stability should therefore permit quantitative insights into membrane association as well as its role in aggregation.

Here we make use of this potential and report on a comprehensive NMR investigation of the effects of lipid charge, bilayer fluidity and αS acetylation on the structural aspects of αS membrane association. We corroborate these insights with molecular dynamics (MD) simulations as well as a series of complementary biophysical measurements to further characterize membrane plasticity, overall affinities as well as binding and aggregation kinetics. Based on this data we correlate structural insights, such as residue specific affinities and competition for accessible membrane surface area, to their potential role in modulating αS aggregation properties. Our study provides insights into (i) the different lipid binding modes of αS to stable planar bilayers of defined lipid quantity and composition, (ii) the effect of membrane plasticity for αS binding, (iii) the modulation of membrane plasticity through αS, and (iv) the connection between binding modes and their effect on αS aggregation. Additionally, it gives an initial estimate of the number of lipid-associated αS molecules that are required to induce/promote nucleation, and allows to develop a basic structural, thermodynamic and kinetic model of the modulation of αS aggregation through its interaction with different membrane surfaces.

Our *in vitro* data help to better understand the molecular determinants of αS-membrane association and point to possible *in vivo* implications in the context of Parkinson’s disease.

## RESULTS

### Effects of phospholipid head-group charge on the αS membrane binding mode

To obtain residue specific insights into the interaction of αS with lipid bilayer nanodiscs (NDs) of various composition, we recorded a series of solution NMR 2D TROSY-HSQC spectra (see Figure 1 – figure supplement 1 for full list of measured samples). The NMR spectrum of αS in the presence of NDs containing only DMPC lipids perfectly overlays with the spectrum of αS in the absence of NDs (Figure 1a-d). This finding provides additional evidence that αS does not interact with non-charged lipid bilayers (Rhoades, et al., 2006), a point that is still controversial in the literature (Davidson, et al., 1998). Additionally, it also shows that αS does not interact with the membrane scaffold protein (MSP), confirming that the effects described in the following are not biased by (unspecific) αS-MSP interactions.

**Figure 1.**
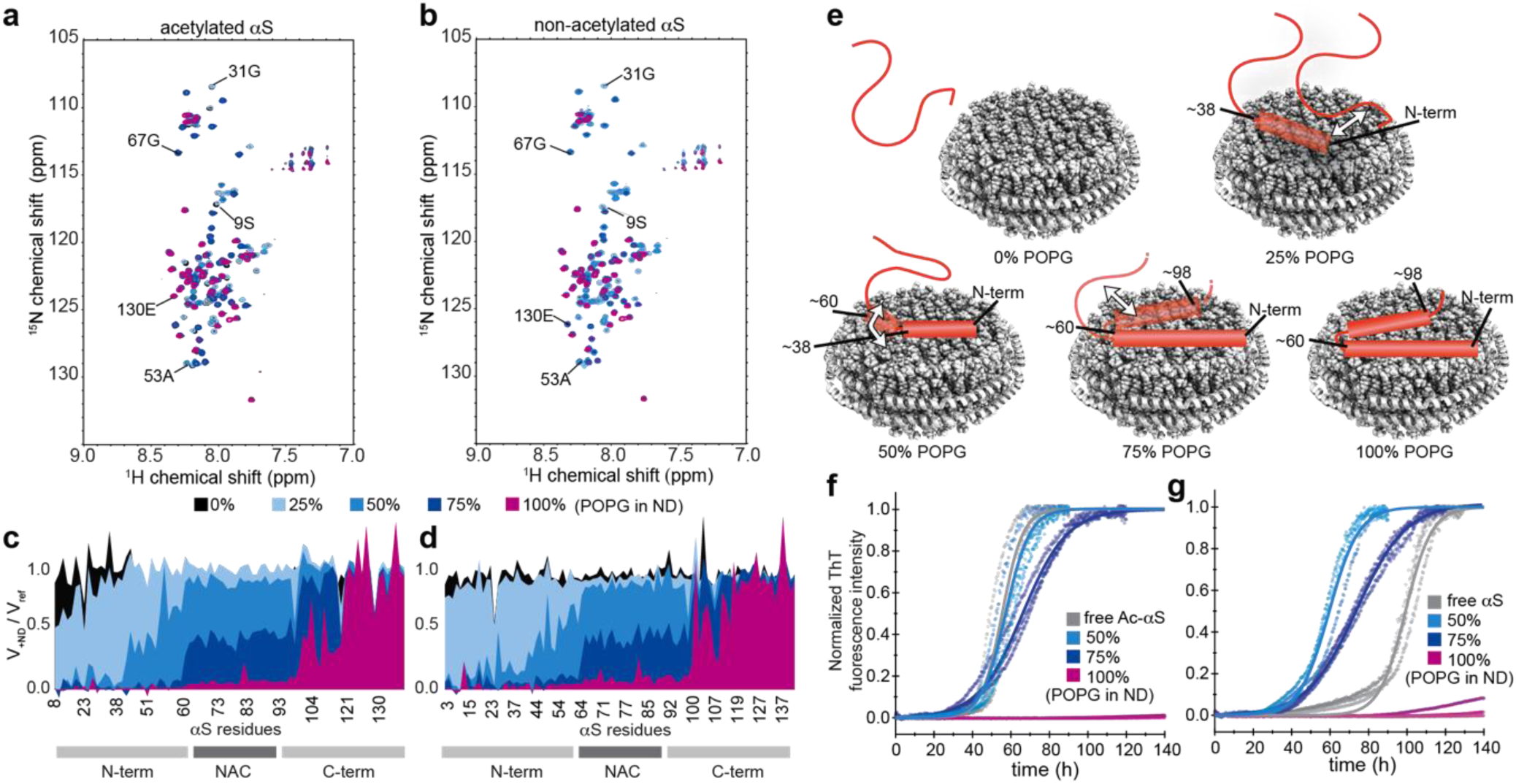
Lipid charge content modulates αS membrane binding modes and different binding modes show different effects on αS aggregation properties. NMR spectra of [^15^N]-αS (50 μM) in the absence or in the presence of 25 μM NDs of different POPG (negative charges) contents. [^15^N-^1^H]-TROSY-HSQC spectra of acetylated (a) and non-acetylated (b) αS. Spectra in the absence (grey) or in the presence of NDs containing 0% POPG (black), 25% POPG (light blue), 50% POPG (blue), 75% POPG (dark blue) and 100% POPG (purple) are displayed. Selected residue assignments corresponding to differently affected parts of αS are indicated. Corresponding NMR attenuation profiles, i.e. the ratio of peak volumes in the presence and absence of NDs, are plotted against αS primary sequence for acetylated (c) and non-acetylated (d) αS. (e) Molecular model visualizing the gradual binding of different parts of αS to NDs with increasing charge content. (f,g) αS aggregation assays (normalized ThT fluorescence) in the absence and presence of NDs with indicated POPG charge content for acetylated (f) and non-acetylated (g) αS (data of triplicate measurements until reaching saturation and their respective fits are shown, color code as in a-d).

In a similar way as reported previously using liposomes (Drescher, et al., 2008, Iyer, et al., 2016), we further tested the influence of increasing amounts of negatively charged lipid head groups on αS membrane association, keeping a molar ratio of one αS molecule per membrane leaflet (Figure 1a-d). Note that lipid ratios and proper mixing of the different lipid types inside the nanodiscs was also observed by NMR spectroscopy (Figure 1 – figure supplement 2a). Our NMR data show a gradually increasing bilayer interaction of αS with increasing lipid charge content, dividing the protein into rather distinct regions with different binding behaviors (Figure 1e). The first region spans the N-terminal residues 1-38, which are already weakly interacting at 25% content of negatively charged lipids and strongly interact at 50% (or higher) charge content. The region of residues 38-60 interacts more gradually at 50% charge content. Amino-acids 60-98, corresponding approximately to the aggregation-prone non-amyloid-β component (NAC region), displays some interactions with membranes containing 75% anionic lipids and strongly interacts at 100% anionic lipid content. The 98-120 region is (partly) affected by 100% net charge content only. Finally, the last 20 C-terminal residues never show any membrane interaction. This data is largely in line with an expected predominantly electrostatic model (the first 60 residues displaying a net positive charge, the last 40 residues a net negative charge, and the NAC region being mostly hydrophobic), as well as the three regions dynamic model reported before using SUVs (Fusco, et al., 2014).

Using Thioflavin T (ThT) fluorescence as a reporter for fibril formation, we also measured aggregation kinetics of αS in the absence and presence of the different ND compositions (Figure 1f-g). These experiments were performed under conditions where αS amyloid fibrils form spontaneously, mainly by interface-driven nucleation and subsequent amplification through fragmentation (non-repellent plates, glass balls, shaking, see below for complementary kinetic assays) and therefore mainly report on the potential interference of nanodiscs with the lipid-independent aggregation pathway of αS (Campioni, et al., 2014, Gaspar, et al., 2017, Rabe, et al., 2013, Vacha, et al., 2014). Interestingly, despite the fact that the NMR data show interaction, the presence of NDs up to an anionic lipid content of 50% does not appear to affect aggregation kinetics (note that the free non-acetylated αS reference (Figure 1g, gray) shows a different behavior, this point will be discussed below). When increasing the negative charge content to 75% the aggregation half-time slightly increases (Figure 1f,g dark blue) and a strong aggregation-inhibiting effect is detected in the presence of NDs with 100% anionic lipids (Figure 1f,g purple).

While the NMR data visualize the modes of αS binding to membranes of different charge contents, the ThT kinetic data allow to directly link these molecular determinants to their effect on αS aggregation. In this respect, one of the most striking connections is that αS interaction with NDs comprising up to 50% negatively charged lipids does not involve the NAC region and that under the same conditions no detectable effect on the aggregation behavior of (acetylated) αS is found in ThT assays. When further increasing the charge density above 50% negatively charged lipids, NMR data show first a partial (75% POPG, Figure 1a-d, dark blue) and then a full (100% POPG, Figure 1a-d, purple) signal attenuation of the NAC region. This correlates well with a slight inhibitory effect of the 75% charged NDs on αS aggregation (Figure 1f,g dark blue) and a very strong inhibitory effect of 100% charged NDs (Figure 1f,g, purple). Our data strongly suggest that for the tested conditions (high anionic lipid content and high lipid-to-αS ratios) membrane association of the NAC region seems to be the dominant factor for protecting αS from aggregation.

### Nanodisc-bound state of αS is predominantly α-helical

It is worth noting that the NMR results described above mainly refer to the decrease in peak intensity as a reporter for interactions, which is in line with the effects seen before using SUVs (Bodner, et al., 2009). While it is clear that SUVs have particle sizes (associated with slow tumbling rates) well above the detection limit of conventional solution NMR techniques, the smaller size of the ND system should, in principle, allow detection of NMR signals, as has been reported before for several ND-bound or ND-integrated proteins (Viegas, et al., 2016). Nonetheless, neither the usage of Transverse Relaxation Optimized Spectroscopy (TROSY) (Pervushin, et al., 1997) with increased signal accumulation (i.e. 10-fold longer as for spectra shown in Figure 1a-b) nor the measurement at increased temperatures (35°C) and the usage of an NMR-optimized smaller membrane scaffold protein (MSP1D1ΔH5) (Hagn, et al., 2013) forming NDs of smaller size and higher tumbling rates, resulted in appearance of a new set of peaks indicative for the bound sate (and the presence of slow exchange processes) or a collective shift of peaks indicative of fast on-off exchange processes (e.g. Figure 3 – figure supplement 1). In theory, three effects may explain this observation and obstruct
detection of the ND-bound residues of αS: (i) the bound-to-free exchange rate is in the order of the NMR time scale (so-called intermediate exchange), (ii) the presence of a non-negligible part of αS protruding out of the ND (namely at least residues 98-140), slowing down molecular tumbling and increasing relaxation leading to line broadening beyond the detection limit, and/or (iii) membrane-bound αS shows a significant amount of plasticity leading to inhomogeneous broadening of the NMR lines. While intermediate exchange can be largely ruled out due to the observed binding kinetics (*vide infra*, Figure 4a,b, t_on_ and t_off_ of around 4 μs·M and 65 s, respectively) it is at this point difficult to further distinguish between slower tumbling and molecular plasticity (or a combination thereof) that interfere with solution NMR detection of the bound conformation.

**Figure 4.**
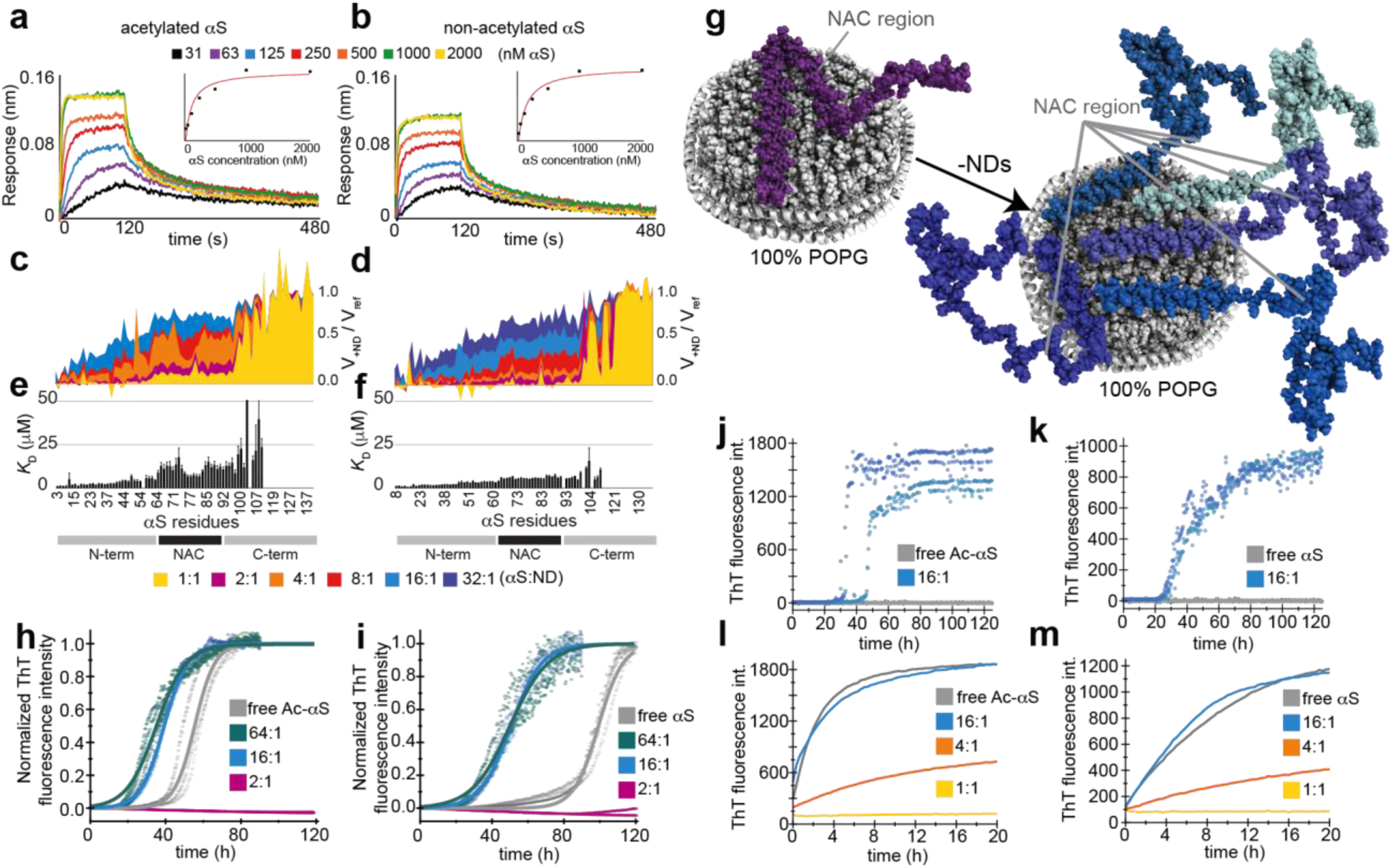
Limited but highly charged lipid surface area can induce an αS binding mode capable of promoting aggregation through induction of primary nucleation. BLI sensorgrams using 100% POPG immobilized NDs and acetylated αS (a) or non-acetylated αS (b) in different concentrations ranging from 31 (black) to 2000 nM (yellow). Corresponding steady-state response plots are shown as inserts. The extracted global affinities (*K*_D_) are about 60 and 100 nM, respectively. NMR attenuation profiles of a titration of 50 μM acetylated (c) or non-acetylated (d) αS with varying concentrations of 100% POPG NDs (αS-to-NDs molar ratios ranging from 32:1 to 1:1, see color code). (e,f) Corresponding residue-specific affinities (*K*_D_) extracted from NMR titration data shown in (c,d), respectively. (g) Model of different αS-lipid binding modes in the presence of excess (upper picture) or limited amount (lower picture) of highly charged NDs (for simplicity only one side of the bilayer is occupied by αS). (h,i) Normalized ThT fluorescence kinetic curves for selected αS-to-ND ratios for acetylated (h) and non-acetylated (i) αS. While high amount of NDs (upper binding mode in g) inhibit aggregation, limited amount of highly charged membrane surface (lower binding mode in g) enhances aggregation. (j,k) Normalized kinetic data in quiescent conditions and absence of preformed seeds at pH 5.3 for acetylated (j) or non-acetylated (k) αS. While under these conditions no aggregation (nucleation) is observed in the absence of NDs (grey) the presence of 16:1 molar ratio of 100% POPG NDs (lower binding mode in g) induces primary nucleation. (l,m) Raw data of quiescent ThT fluorescence aggregation assay in the presence of 2.5% preformed seeds for acetylated (l) or non-acetylated (m) αS for different molar ratios (see color code) of 100% POPG NDs.

In order to still gain insight into the conformation of αS bound to NDs, we used magic angle spinning (MAS) solid-state NMR which is not subject to size effects. Moreover, we took advantage of the very low temperatures (100 K) used in Dynamic Nuclear Polarization (DNP) to additionally eliminate exchange processes, as well as to increase the sensitivity of the experiment. To avoid problems of signal overlap arising from severe inhomogeneous line broadening often seen in this range of temperatures (Siemer, et al., 2012), we used a sparse isotope labelling scheme (Hong and Jakes, 1999), leading to the simplification of ^13^C-^13^C spectra to secondary structure sensitive Cα-Cβ cross-correlations of valines (and leucine Cβ-Cγ). Notably according to the αS primary sequence (Figure 2, top) and our solution NMR observations (Fig. 1a-b), 95% of the valine residues (i.e. 18 out of the 19) should be membrane-bound at the used charge content and αS-to-ND ratio. While in the absence of NDs the DNP ^13^C-^13^C spectrum (Figure 2, black) shows a continuous distribution of the Valine Cα-Cβ cross peaks reflecting the carbon chemical shifts of the allowed Ramachandran space (expected for an intrinsically disordered protein such as αS), a very strong peak shift to a defined chemical shift range typical for α-helical structure is visible after addition of NDs (Figure 2b, red). The DNP data thus show that αS binds the ND lipid surface in α-helical conformation corroborating previous studies using CD and vesicles, solution NMR and detergents micelles and solid-state NMR and SUVs (Fusco, et al., 2014, Galvagnion, et al., 2015, Ulmer, et al., 2005).

**Figure 2.**
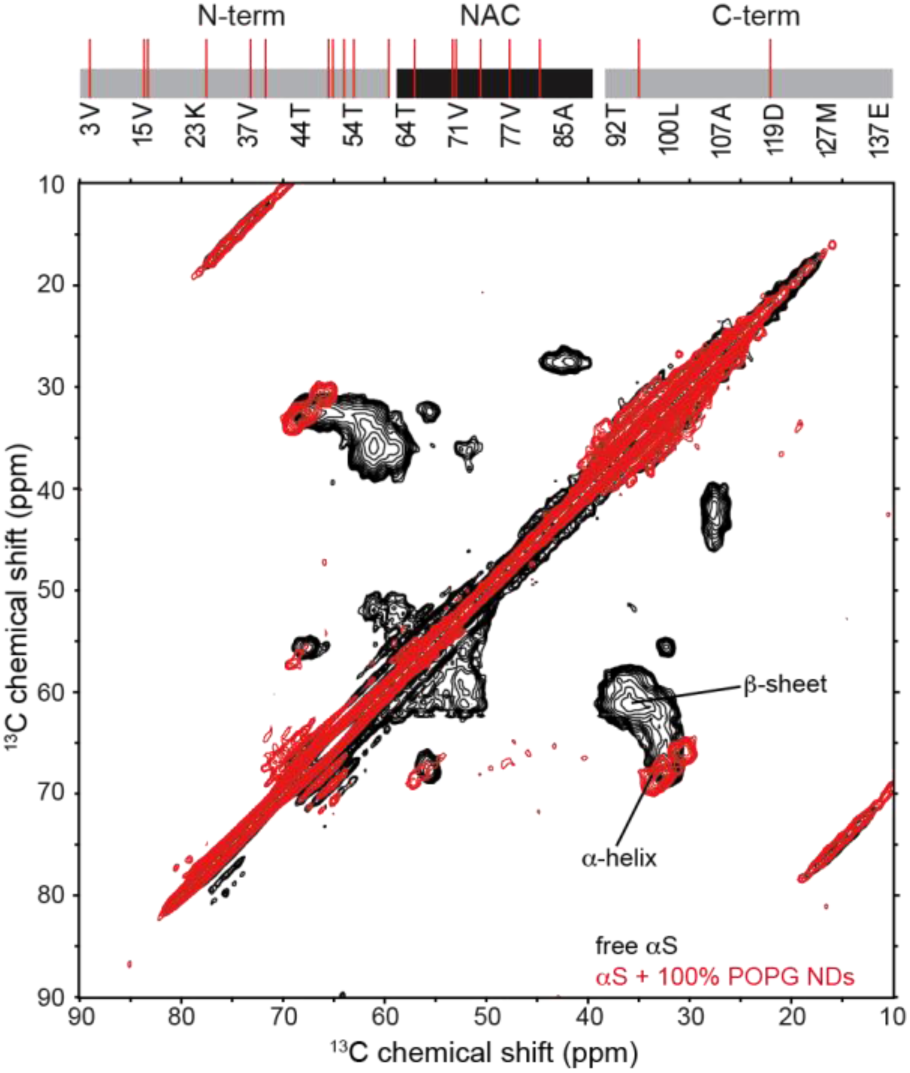
Nanodiscs binding induces α-helical structure in αS. [^13^C-^13^C]-Proton Driven Spin Diffusion MAS-DNP spectra of free αS in frozen solution (black) and when bound to NDs with 100% POPG lipids (red). Selective isotope labelling was used to specifically monitor Valine Cα-Cβ chemical shift distribution. Peaks position indicative of the β-sheet and α-helical secondary structure are labelled. Occurrence of Valine residues in the αS sequence is shown on top (18 out of 19 Valines are present in the expected binding site of the used ND).

### Membrane plasticity and αS-membrane interaction: a two-way street?

While Figure 1 shows the effect of gradually increasing the level of POPG in DMPC lipids it should be noted that both lipids do not only differ in their head group charge but also in their hydrocarbon chain length and number of unsaturated bonds, i.e. 16:0-18:1 PG and 14:0 PC for POPG and DMPC, respectively (nomenclature refers to ‘number of carbons in the fatty acid’: ’number of unsaturations’). Both of these features will affect the fluidity and phase transition temperature T_m_ of the lipid bilayer. In general, native membranes are heterogeneous mixtures of numerous different lipids and proteins, both of which can strongly influence lipid phase properties. In this respect it was also suggested before that the MSP-lipid interactions may not only be seen as an artificial border of the bilayer, but may partly mimic protein-lipid interaction occurring in native membranes (Alvarez, et al., 2010). While the usage of additional transmembrane proteins is rather restricted due to the limited bilayer area of the NDs, the usage of different heterogeneous lipid mixtures may generate a suitable mimetic for rather heterogeneous physiological membranes.

To further investigate the effect of different lipid properties we recorded additional NMR spectra of αS in the presence of NDs containing different lipids and lipid mixtures. The data recorded with 100% POPC NDs do not show interaction (Figure 3a, grey), comparable to 100% DMPC NDs (Figure 1a-d, black) and demonstrates that in the absence of negative net charges on the head groups, properties such as an increased chain length and unsaturation, do not induce a significantly stronger interaction. We also used lipid mixtures where charged and uncharged lipids have the same fatty acid chains (i.e. POPC/POPG and DMPG/DMPC) or mixtures where the position of negative charge and hydrocarbon chain is swapped (i.e. DMPG/POPC vs. POPG/DMPC). In these mixtures, we kept the overall net charge content constant at 50%. Our data show that the heterogeneous mixtures DMPG/POPC and POPG/DMPC behave very similarly (Figure 3a, turquoise and dark blue, respectively) suggesting that the position of charge with respect to chain length and unsaturation is not critical in the tested conditions. Also, when using a more homogenous mix of POPG/POPC, a similar pattern is found (Figure 3a, yellow), suggesting that ‘surface roughness’ of the membrane, as potentially introduced by the heterogeneous lipid chain lengths, does not significantly change the αS-membrane interaction.

**Figure 3.**
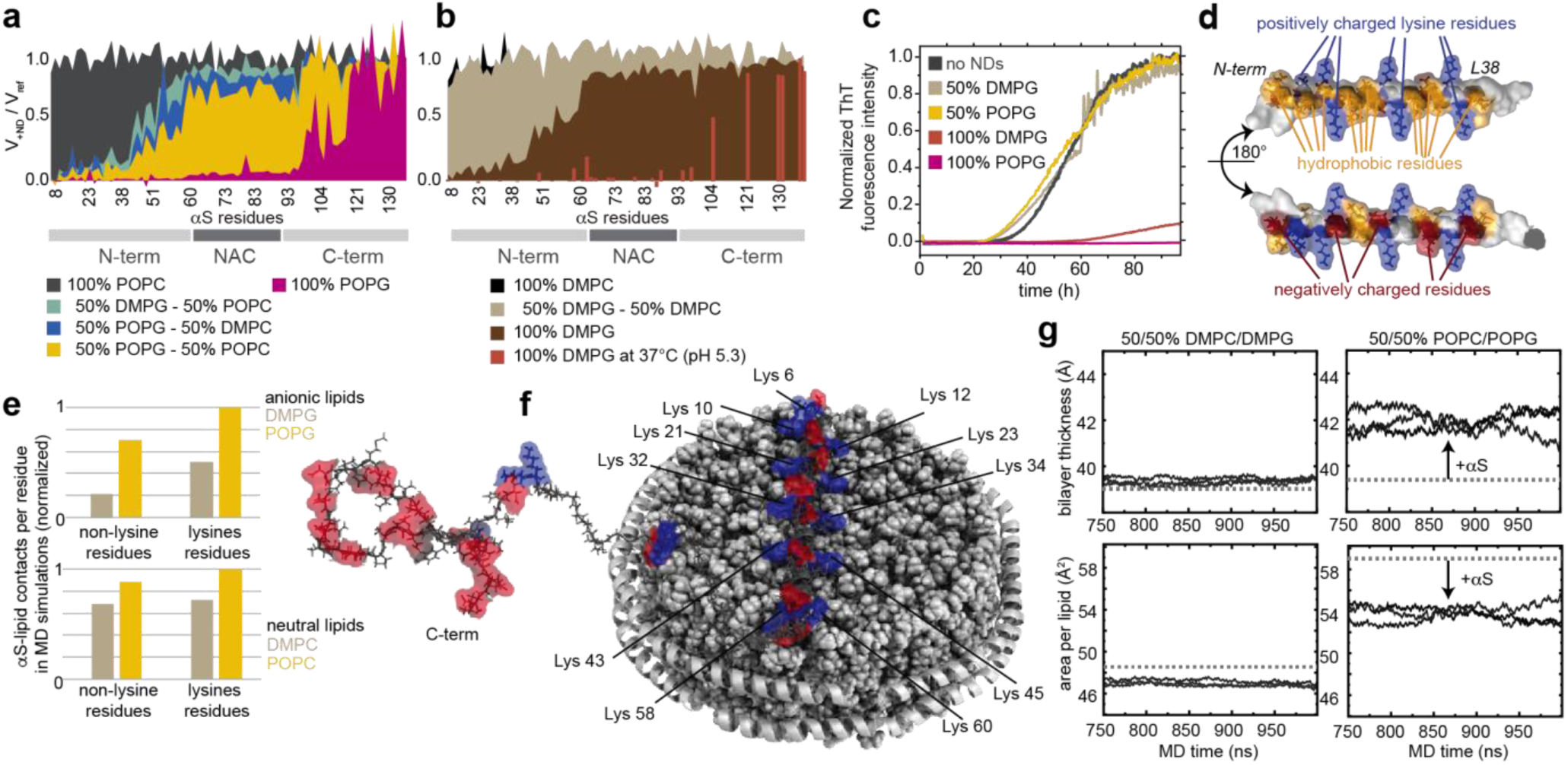
αS-lipid interaction is favored by, and potentially modulates, membrane plasticity. αS NMR attenuation profiles in the presence of NDs with indicated lipid composition (molar ratio 2:1, αS-to-NDs) for bilayers that are in a more fluid (**a**) or gel phase (**b**) at 10°C. Note that all NMR data, except the indicated 100% DMPG measurement (b, red bars), are recorded at 10°C. (**c**) Corresponding ThT aggregation assays for selected conditions (color code as in (a,b), for better visibility only the mean values of triplicate measurements are shown). (**d**) αS structural model of key lipid binding mode (see text for more details). Periodically and symmetrically appearing lysine residues (blue) form a positively charged ‘grid’. (**e**) αS-to-lipid contacts (<4 Å) per residue as occurring during the time course of MD simulations. Normalized values for interactions of either lysine or all other residues with anionic lipids (upper diagram) or neutral lipids (lower diagram) for gel/fluid-phase membranes (beige/yellow bars) are shown, respectively. (**f**) Model of αS-ND interaction, lysine (blue) and negatively charged residues (red) are highlighted. (**g**) Effect of αS interaction in MD simulations on the two different membrane systems. Bilayer thickness (upper panels) and area per lipid (lower panels) in the presence (solid lines, three independent simulations) and in the absence of αS (dashed line, one simulation) are shown.

Interestingly, when using a homogeneous mix with fully saturated lipids (DMPC/DMPG), albeit having the same net charge content of 50%, a very different pattern in line with no significant interaction with αS, is visible (Figure 3b, beige). Notably, at the temperature of the NMR experiments (10 °C) the bilayer formed by DM-lipids is in the gel phase (T_m_ around 28 °C, Figure 1 – figure supplement 2d). This data is in line with previous data on SUVs that identified an important role of the lipid phase for αS-lipid interaction (Galvagnion, et al., 2016). When increasing the charge content to 100%, but remaining in the gel phase (100% DMPG, Figure 3b, brown) the first 60 residues of αS show a clear interaction with the membrane resembling a binding mode that is found for 50% charge content in the fluid phase (e.g. Figure 3a, yellow). While increasing the temperature for the NMR measurements above Tm leads to a previously observed loss of NMR signals due to amide-water exchange processes for most residues relevant for the interaction (Figure 3 – figure supplement 1c-d), lowering the pH from 7.4 to 5.3 (Figure 3 – figure supplement 1e-f) can counter this effect and allows to confirm that when measuring the interactions to DMPG lipids in the fluid phase, a much larger binding interface is found (Figure 3b, brown-red), comparable to the binding mode found for 100% POPG (Figure 3a, purple). Consequently, when the lipid charge density is high enough, the lipid phase state and related bilayer fluidity in a stable planar bilayer plays an important role in αS association. Taken together, these data on αS-lipid association can be summarized as follows: (i) unsaturations in the hydrocarbon chains, leading to increased membrane fluidity, are not sufficient to induce binding, (ii) the presence of heterogeneity in fatty acid chains and the combination of charge and unsaturation on the same lipid molecule are not critical and (iii) in addition to charge, a lipid phase state that introduces an increased membrane fluidity is important for binding.

We also performed ThT aggregation assays with NDs containing selected lipid mixtures as investigated by NMR, again under conditions in which spontaneous aggregation of αS is observed in the absence of lipids. All mixtures that contain 50% negatively charged lipids, independently of tail heterogeneity or charge position show consistently unaffected aggregation behavior (Figure 3c). However, αS aggregation is drastically impeded in the presence of 100% DMPG NDs (Figure 3d, red-brown). Since aggregation assays were measured at 37 °C (i.e. above the DMPG phase transition) this is well in line with the corresponding NMR attenuation profile (Figure 3c, red-brown) suggesting that under this conditions αS is in a lipid binding mode that involves the NAC region and thus inhibits aggregation.

From the evaluation of the different membrane interaction modes of αS with NDs observed in this study, it is clear that in particular the initial about 38 residues comprise a central lipid binding motif. Based on previously reported solid-state NMR data (Fusco, et al., 2014) and CD data (Galvagnion, et al., 2015, Jo, et al., 2000) of vesicle preparations as well as the DNP solid-state NMR data of αS in the presence of NDs that we report here, it is also clear that the lipid interacting residues of αS form a helical secondary structure. Analysis of the primary sequence and secondary structure of this interacting region highlights key features of its architecture which is highly suitable for interactions with charged lipid (bi)layers (Figure 3d) (Davidson, et al., 1998). These features include the very exposed and symmetric distribution of positively charged lysine side chains (Figure 3d, blue), the occurrence of hydrophobic residues on one side of the helix (Figure 3d, yellow) and the distribution of negative charges on the opposite side (Figure 3d, red). Based on this architecture it is tempting to speculate that the positively charged residues will interact with negatively charged lipid head groups, the hydrophobic residues will be oriented towards the hydrophobic lipid chains and the negatively charged residues will be oriented towards the solvent (and compensate the net charge of the protein). Noteworthy these protein features are again found in the next binding region (residue 39-60). In this picture, it would be likely that the lipids as well as the lysine side chains (partly) rearrange, from their ‘unbound’ conformation, to ideally accommodate electrostatic interactions in the bound conformation. This rearrangement may be favored by a more fluid lipid phase, which would explain the lower interaction found for NDs with DMPG lipids below the phase transition.

To test this hypothesis, we performed molecular dynamics (MD) simulations of αS-membrane interactions. Our simulations focus on the first 61 residues of αS and its interactions with membranes formed by a mixture of either 50% POPG - 50% POPC lipids in the fluid phase or a 50% DMPG - 50% DMPC mix in the gel phase (see methods for more details). Indeed, the MD data confirm that the lysine residues play a key role in the membrane interaction, as e.g. visible by promoting considerably more contacts to anionic lipids than other residues during the time course of the simulations (Figure 3e, upper diagram). Additionally, the MD data also show a generally stronger interaction of αS with the anionic lipids in the fluid membranes (POPG) as compared to the gel-phase membranes (DMPG) (Figure 3e, yellow vs. beige). Noteworthy, these effects are much less pronounced for contacts to the neutral lipids (Figure 3e, lower diagram). These findings correlate well with the effects of lipid charge and membrane plasticity seen in the NMR and aggregation assays.

Interestingly, the MD data also report on the effect of αS interaction from the lipid point of view. According to this data, the already well ordered DMPC/PG-lipids (gel phase) experience only very small effects due to the presence of αS. On the other hand, the less ordered POPC/PG-lipids (fluid phase) are strongly affected. Here the presence of αS induces a considerably more ordered lipid state as evident by an increased bilayer thickness, reduced surface area per lipid and increased order parameters for the hydrocarbon chains (Figure 3g and Figure 3 – figure supplement 3). Generally, the MD data suggest that αS-membrane interaction is (initially) facilitated by increased membrane plasticity (e.g. more contacts for fluid phase). The increased plasticity may facilitate reorientations of the lipids to support favorable interactions with αS. This view is supported by our MD data showing significantly higher occurrence of short lysine-lipid distances, which would allow lipid-mediated salt bridges, in the simulations with more fluid membranes (Figure 3 – figure supplement 3). These interactions may consequently confine lipids and lead to a reduced membrane plasticity. The latter is in line with recent experimental data showing that αS binding can increase lipid packing (Iyer, et al., 2014) an effect that has also been suggested to play a role in αS function as chaperone for SNARE-mediated vesicle fusion (Burre, et al., 2010).

### NDs can simultaneously interact with multiple αS proteins facilitating formation of aggregation seeds

While our NMR data clearly reveal binding modes with different contributions of the αS primary sequence, we were also interested in the overall affinity of the protein for NDs. We therefore measured interaction kinetics using bio-layer interferometry (BLI) with immobilized NDs of different charge content. In line with the NMR data, no αS binding was detected when NDs containing 100% DMPC were immobilized (data not shown). In the case where NDs with 100% anionic lipid content were immobilized, a clear response upon addition of different αS concentrations was observed (Figure 4 a,b) and dissociation constants *K*_D_ in the order of 60-100 nM (one αS to one ND) were calculated. In addition, kinetic information could be extracted which shows a tight binding with a fast association and a slow off-rate of approximately 10^−2^ s^−1^.

In order to obtain residue-specific insights into the membrane affinity of αS, we additionally conducted NMR titration experiments using 100% negatively charged NDs (Figure 4c-f). In general, dissociation constants can be extracted from NMR titrations attenuation profiles by fitting the concentration dependency of the attenuation with a single exponential decay for each resolved peak (corresponding to one assigned residue) (Figure 4e-f). Our data reveal differential membrane affinities for different regions of the αS primary sequence. The regions with differential affinities largely overlap with the regions of the different binding modes identified before, i.e. four distinct regions of decreasing affinity range (1-38, 39-60, 61-98, 99-140).

Importantly, it appears that one ND with 100% negatively charged lipids can simultaneously interact with up to 16 αS molecules (8 per bilayer side) in the course of the NMR time scale, as seen from the almost complete disappearance of the signals of the very N-terminal residues (Figure 4e-f, light blue). Moreover, while these first residues are interacting with the membrane, independently of the number of αS molecules bound to one ND, the binding of the NAC region is strongly modulated by the number of bound αS molecules. This suggests that, due to the higher lipid affinity of the N-terminal region as compared to the NAC region, the free energy of the system is minimized by favoring N-terminal interactions in cases where the accessible membrane surface is limited. Note that due to the geometry of the used nanodiscs up to five αS molecules can simultaneously bind with a 38-residue long α-helix (first binding mode) to one side of one ND. If 8 molecules are accommodated together on the surface, (on average) a 23-residue long helix per monomer could be formed.

To characterize the effect of the accessible membrane surface area (as given by the αS-to-ND ratio) and the resulting stoichiometry on αS aggregation behavior, we measured ThT aggregation kinetics on samples with different αS-to-ND ratios ranging from 2:1 to 64:1 by varying the ND concentration at constant αS concentrations (Figure 4h-i). Interestingly, a higher ratio of αS-to-ND leads to a prominent decrease in aggregation lag-times when using 100% POPG NDs (Figure 4h,i, blue and cyan). These data show, in line with previously reported behavior on SUVs (Comellas, et al., 2012, Galvagnion, et al., 2015, Zhao, et al., 2004), that under specific conditions lipid bilayers can accelerate the fibrillation process. Here, these conditions represent a limited membrane surface area with a high charge density. According to our NMR data this will introduce an αS-lipid binding mode that brings several αS molecules with exposed NAC regions in close proximity.

Noteworthy, it has been reported in the case of SUVs (Galvagnion, et al., 2015, Hellstrand, et al., 2013) that lipids can be incorporated in fibrils and change their morphology. Nonetheless, in the case of the more stable nanodiscs this is not expected, indeed AFM images of αS fibrils formed in the absence or presence of NDs do not show different morphology (Figure 4 – figure supplement 1) and the size of nanodiscs does not change after interaction with αS (as seen from SEC, Figure 1 – figure supplement 2). This indicates that a perturbation of the aggregation pathway by lipid incorporation is very unlikely in our case.

We additionally carried out the same BLI measurements, NMR titrations and ThT assays for ND containing only 50% POPG lipids. For these NDs no clear signature of binding could be obtained in the BLI measurements (data not shown), suggesting a weak affinity and/or too fast off rates to allow detection via BLI. This is in line with SEC profiles that also point to a more transient interaction (Figure 1 – figure supplement 2).

NMR titrations, however, show clear concentration dependent attenuation profiles that allow the calculation of residue specific affinities (Figure 5a-d). Noteworthy, the affinities for the αS residues in the first binding region (residues 1-38) are comparable to the values obtained for 100% charged NDs (Figure 4e-f). In contrast, for the following binding regions much lower affinities are found (at the edge of detection for residues 39-60, and no interaction for residues > 60), including the absence of interactions of the NAC region. In line with an exposed NAC region the ThT data for NDs with 50% negatively charged lipids at low αS-to-ND ratios are consistently showing no effect on aggregation half-times. The data at higher ratios are less reproducible and show a slight tendency to prolonged half-times (Figure 5e-f). At this point, it is not clear whether this feature has mechanistic relevance or is just an artefact caused by the limited reproducibility of this condition.

**Figure 5.**
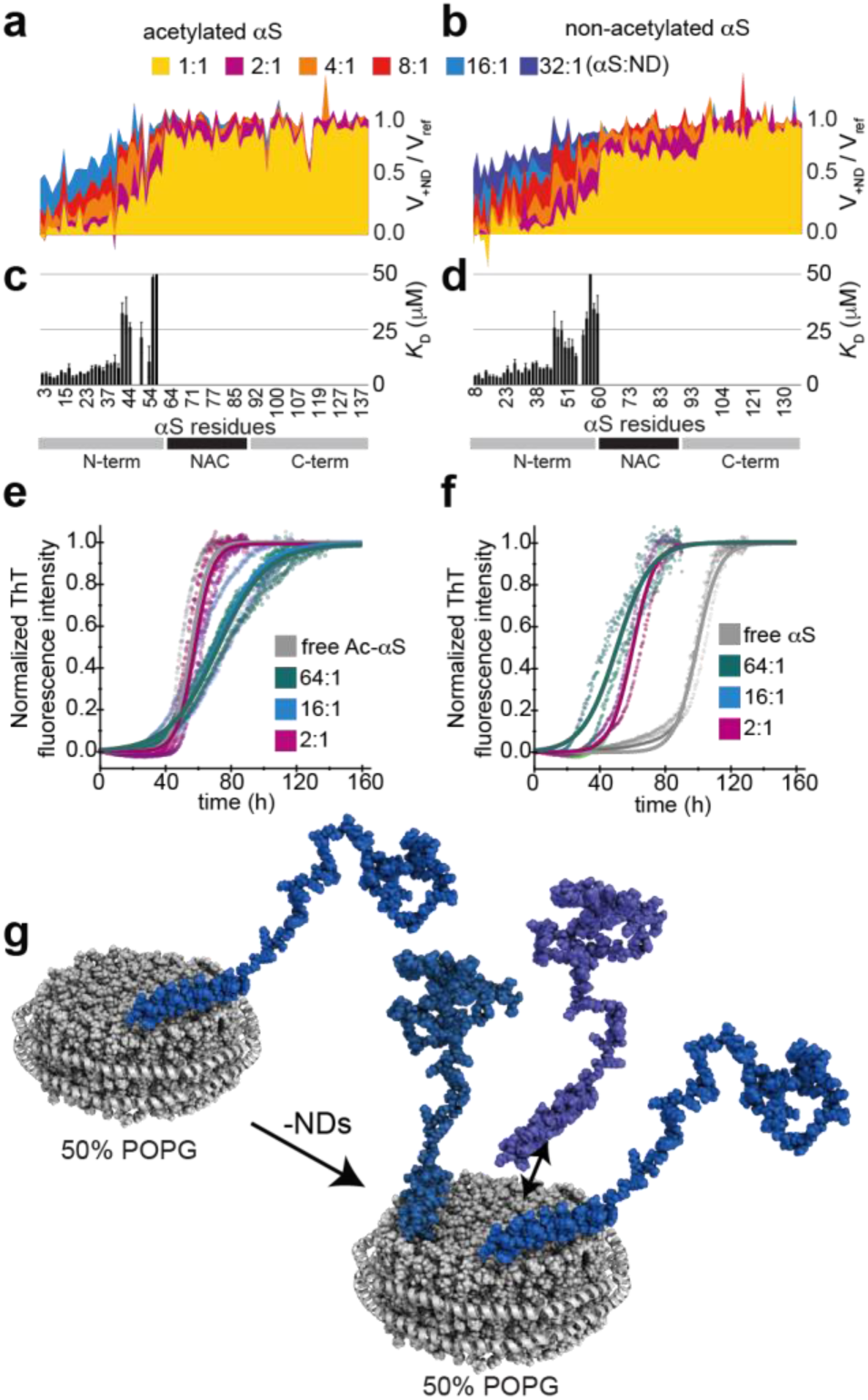
Moderately charged membrane surfaces interact more transiently with αS and do not induce primary nucleation. NMR attenuation profiles of a titration of 50 μM acetylated (a) or non-acetylated (b) αS with varying concentrations of 50% POPG NDs (αS-to-NDs molar ratios ranging from 32:1 to 1:1, see color code). (c,d) Corresponding residue-specific affinities (*K*_D_) extracted from NMR titration data shown in (a,b), respectively. (e,f) Normalized ThT fluorescence kinetic curves for selected αS-to-ND ratios for acetylated (e) and non-acetylated (f) αS. (g) Model of αS-lipid binding modes in the presence of excess (upper picture) or limited amount (lower picture) of moderately charged NDs (for simplicity only one side of the bilayer is occupied by αS).

### The molecular, thermodynamic and kinetic determinants of membrane-modulated αS aggregation

It has recently been shown in a systematic study that under appropriate conditions, which minimize the intrinsic nucleation rate (quiescent conditions and protein repellent plate surfaces), lipid bilayers in the form of small unilamellar vesicles can largely accelerate the nucleation of αS amyloid fibrils (Galvagnion, et al., 2015). In order to determine whether NDs can have a similarly accelerating effect, we performed ThT aggregation experiments under similar conditions, i.e. where no αS aggregation should be detected in the absence of lipids (see methods for more details). These experiments were performed at mildly acidic pH (5.3), as it was recently shown that under these conditions, αS amyloid fibrils can amplify autocatalytically through surface-catalyzed secondary nucleation (Buell, et al., 2014, Gaspar, et al., 2017). This should in principle enable even very low primary nucleation rates to be detected through autocatalytic amplification. Membrane binding behavior of αS at these buffer conditions was also checked by NMR for 100% POPG NDs and comprises the same region (residues 1-96 at low αS-to-ND ratios) as seen for neutral pH (Figure 3-figure supplement 1c,d).

As expected, the ThT assay under the quiescent conditions does not show any aggregation in the absence of NDs (Figure 4j,k grey). However, the samples with 100% POPG NDs (in a αS-to-ND ratio of 16:1) do display aggregation, indicating that this type of membrane is indeed enhancing primary nucleation to a degree sufficient that the subsequent secondary nucleation leads to detectable quantities of amyloid fibrils (Figure 4j,k blue). Interestingly, since our data also allow an estimation of the total number of αS monomers that are brought in close proximity due to their interaction with the same ND, this result may also provide a first approximation of the number of αS monomers needed for the formation of a nucleus. Our data suggest that this ‘minimal critical nucleation number’ has an upper limit of around 8 αS molecules.

In order to further disentangle the effect of NDs on the various individual steps in the αS aggregation pathway, we next designed strongly seeded aggregation assays under quiescent conditions at neutral pH, where fibril elongation is the only process that occurs at a significant rate. Seeding experiments carried out in the presence of 100% charged (POPG) NDs show a clear reduction in elongation rate with increasing ND concentration (Figure 4l,m). While at high molar excess of αS (16:1, αS-to-ND), according to our NMR data, nearly all αS monomers should interact through their most N-terminal regions with the NDs, no significant effect on the elongation rate was observed (Figure 4l,m blue). At molar ratios of 4:1 (Figure 4l,m orange) a decrease in elongation is observed, however at this ratio our NMR data clearly indicate that all αS monomers are interacting with the NDs. Nevertheless, in both cases (molar ratios of 16:1 and 4:1) larger fractions of the monomer population should still have accessible NAC regions, which may explain their ability to participate in the fibril elongation process. However, the limited interaction surface may also influence the dynamic nature of membrane association leading to a certain population of free monomeric αS at any given time.

Unlike for 100% POPG NDs and in line with the previously discussed moderate effects of 50% POPG NDs on the overall aggregation process (Figure 1f,g blue, Figure 3d yellow and beige, Figure 5e,f), we did not observe accelerated αS nucleation in the presence of NDs with 50% POPG nor a clear perturbation of fibril elongation in seeded experiments (Figure 5 – figure supplement 1c-i). This data is difficult to explain given that elongation is, in all cases of amyloid formation, responsible for the generation of the bulk of fibril mass. Hence, its inhibition should slow down the overall aggregation kinetic, also under non-seeded conditions. Noteworthy, we observe that ThT signal intensity can be strongly affected by the presence of NDs and that the absolute ThT intensity does not correlate with absolute fibril mass when comparing data recorded in the absence or presence of NDs (as seen from SDS-PAGE, Figure 5 – figure supplement 1e). Although our NMR data show that these NDs do not interact with the αS, the corresponding ThT signal (of the identical samples) show very different intensities (Figure 5 – figure supplement 1f). It is therefore likely that the ThT (unspecifically) interacts with NDs, leading to an overall decrease in ThT intensities. We therefore normalized most ThT assays. Noteworthy, this effect is less pronounced for ND with 100% POPG (as e.g. visible in the data in Figure 4m,l), which we attribute to the high coverage of the lipid surface by αS molecules that may reduce the unspecific ThT-ND interactions. In line with our other data recorded using 50% POPG NDs, albeit significantly reduced ThT sensitivity, only very moderate effects of the NDs on the fibril elongation process are visible after normalizing ThT intensities in the seeded aggregation assays.

### N-terminal acetylation has moderate effect on membrane association and leads to different behavior in aggregation assays

As visible in Figures 1 and 3-5, in addition to using the N-terminally acetylated form of αS, which represents the native post-translational modification of αS and is known to be relevant for membrane association (Nemani, et al., 2010, Theillet, et al., 2016), we also recorded most experiments with the non-acetylated form of αS. In line with previous findings (Maltsev, et al., 2012), N-terminal acetylation leads to clear chemical shift perturbations in the NMR spectra for the first 10 residues of αS (Figure 1a,b).

Overall, most of the above discussed features of αS membrane interaction are rather similar in acetylated and non-acetylated αS, however there are a number of distinct differences. For instance, the peaks which are already shifted in free αS due to the acetylation are also the ones that are affected most by the presence of nanodiscs with low amount of charges (close to physiological concentration). Our data (Figure 1a-d) show a rather small but significant increase in the membrane association of the first 15 residues due to the N-terminal acetylation, which is in line with previous observation using SUVs (Bartels, et al., 2014, Dikiy and Eliezer, 2014). αS acetylation is known to increase N-terminal helix propensity (Kang, et al., 2013, Maltsev, et al., 2012), which may facilitate formation of the initial binding mode and be of significance for naturally occurring processes.

A similar effect is also seen for the global binding as determined by BLI, which shows a (slightly) higher membrane affinity of the acetylated (Figure 4a, *K_D_* of 60 nM) as for the non-acetylated αS construct (Figure 4b, *K_D_* of 100 nM). When looking deeper into the NMR titration data, it appears that another effect takes place, namely a slightly increased membrane affinity of the NAC region for the non-acetylated αS NAC region (Figure 4e *vs*. f and Figure 5c vs. d). At current stage, it is not easy to explain why a modification at the N-terminus will affect the lipid interaction of a protein region that is sequentially separated by roughly 60 residues.

Such a behavior could however either be related to intermolecular interactions and/or long range intramolecular interactions (in a ‘horseshoe’-conformation) that may or may not be artificially introduced by the limited surface area of the NDs.

Strikingly, the reference kinetic curve of non-acetylated αS reproducibly shows under the applied conditions a strongly delayed aggregation as compared to the acetylated reference. In the setup used, primary nucleation processes are likely to happen at the air-water or plate-water interface (Campioni, et al., 2014, Vacha, et al., 2014), thus a lower hydrophobic propensity of non-acetylated αS could explain this effect. While this may be the dominant process in the absence of lipids, it may not be the case anymore in the presence of NDs (Galvagnion, et al., 2015), either because NDs shield these interfaces or because nucleation happens primarily at the membrane surface. The much lower differences due to acetylation state in the presence of NDs fit this explanation, as well as additional tests we ran using different types of plates (data not shown). Higher order processes, namely different fragmentation behaviors, can however not be excluded.

It appears that the biggest effect of acetylation is related to assay parameters that are normally not the matter of interest, which nevertheless may be important for future studies (Iyer, et al., 2016). Still the results from systematic measurement of the effects of N-terminal acetylation via different methods point to subtle changes in membrane interaction in respect to NAC region specific affinities at high lipid charge densities as well as to N-terminal binding at a lipid charge density comparable with the overall composition found for membranes in e.g. synaptic vesicles (Fusco, et al., 2014). Hence, both effects may be of particular importance under physiological conditions.

## DISCUSSION

Overall our data demonstrate that the nanodisc system allows to study the interaction of αS with stable, planar membranes in a quantitative and site-resolved way. It also provides insights into the correlation between the identified membrane binding modes as well as binding kinetics and their consequences for αS amyloid fibril formation, both in respect to the nucleation and elongation process. In summary, our data show that (i) residue specific αS-membrane affinities are rather similar for the N-terminal αS region for 100% and 50% negatively charged NDs, (ii) for 100% anionic lipids the αS can adopt a substantially expanded binding mode as compared to 50% anionic lipids content, leading to considerably higher global affinities, (iii) the exchange rate between free αS in solution and membrane-bound αS is rather slow in the 100% charged case and likely to be rapid in the 50% case, (iv) region-specific membrane affinities (especially the NAC region) are correlated with aggregation properties, (v) with sufficient excess of lipids and sufficient charge density, NDs can inhibit primary nucleation as well as fibril elongation by sequestering monomers out of solution, (vi) competition of αS monomers for highly charged lipid surface generates a membrane-bound αS conformation that can induce primary nucleation, and (vii) the number of αS monomers that are brought together on one ND and which can promote amyloid fibril nucleation is in the order of 8 αS molecules. Figure 6 summarizes these molecular, thermodynamic and kinetic determinants of membrane-modulated αS aggregation.

**Figure 6.**
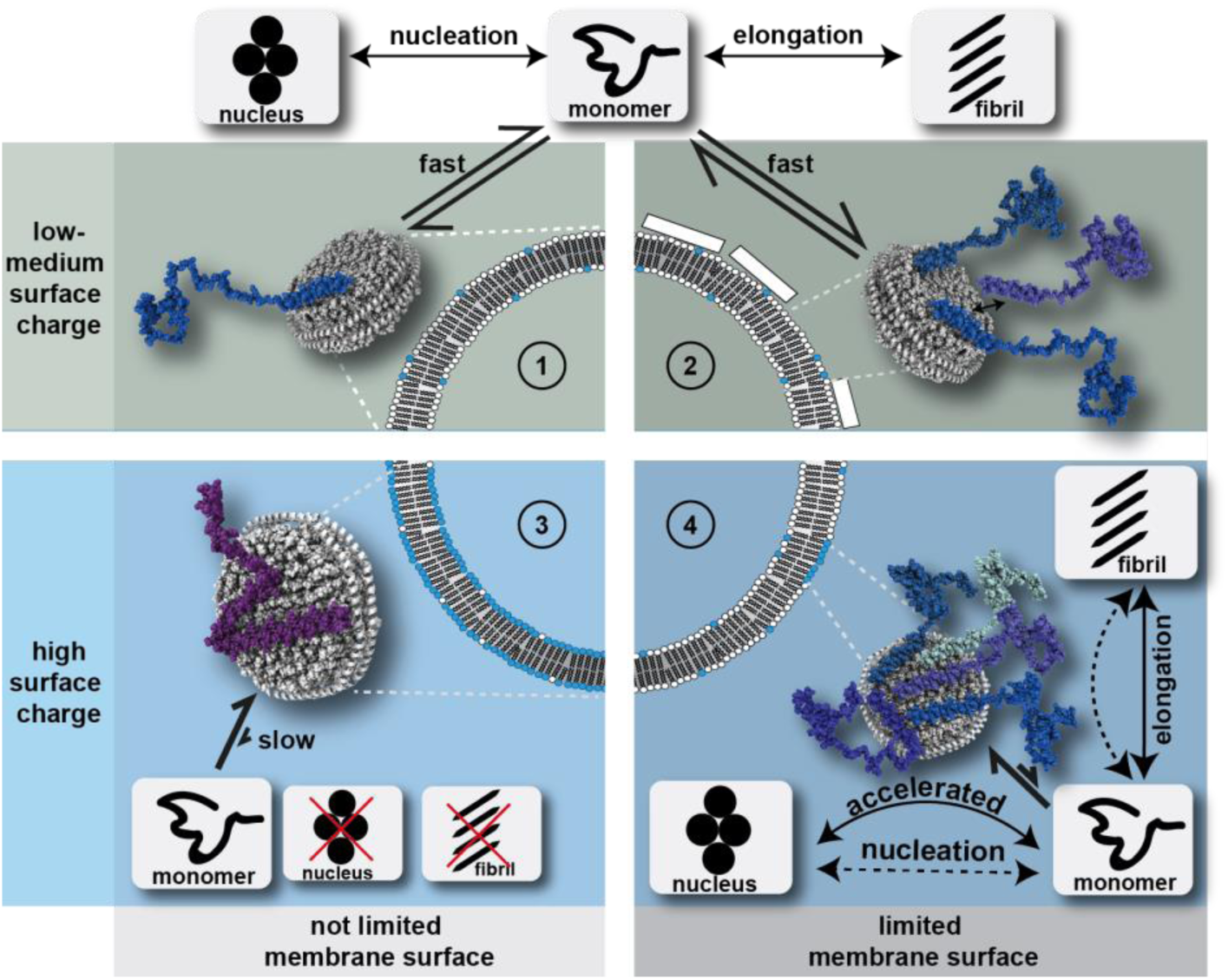
Model of the influence of different NDs on the αS aggregation pathways and its potential implication in the context of cell/vesicle membranes. Four scenarios are depicted according to their membrane surface charge and accessibility, respectively. In cases where membranes/NDs with only low charge densities are present (scenario 1 and 2) αS interacts with its N-terminal residues and most likely forms a fast exchanging equilibrium between soluble and membrane associated αS monomers. This equilibrium does not seem to strongly interfere with the slow process of αS nucleation, it may however (slightly) decrease the pool of free monomers available for fibril formation. Notably these conditions are more likely to better resemble the average charge densities found in physiological membranes. However, specific, abnormal and/or stochastic processes may also lead to highly charged membrane surfaces with limited (scenario 4) or not limited surface access (scenario 3). αS will strongly interact with the latter in a binding mode that will largely inhibit both, αS nucleation and fibril elongation. In cases where several αS monomers compete for a limited highly charged membrane surface area (scenario 4), the amyloid fibril nucleation process can be accelerated, most likely due to an αS binding mode that brings exposed NAC regions of several αS monomers in close proximity. Under these conditions the fibril elongation rate is largely unperturbed, probably due to sufficient monomers with limited membrane association and/or due to induction of higher order processes.

While our *in vitro* data show strongest effects at lipid charge densities well above the average lipid compositions of native membranes, the membrane composition may vary locally in physiological membranes. Due to the normally found high lipid diffusion rates, clusters of higher negative charges may form spontaneously and/or be induced by an initially transient αS interaction. For the latter, the N-terminal acetylation may play an important role, since it increases membrane interaction at average native lipid charge densities.

Our data suggest that clusters of around 60-80 negatively charged lipids suffice to form a strong interaction (this may however not be the lower limit). Sporadically formed lipid charged clusters could also induce a competition of several αS monomers for the accessible surface area. Our data show that due to the different residue-specific membrane affinities this will generate a binding mode that, once the rather low αS critical oligomerization number is reached, can act as an aggregation seed. Such a scenario could promote the initial step of primary nucleation in the pathogenesis of Parkinson’s disease and is in line with recent *in vivo* findings suggesting that shielding αS from membrane interactions can inhibit initial steps of amyloid fibril formation including the formation of cell toxic species (Perni, et al., 2017).

## MATERIALS AND METHODS

### αS and N-terminally acetylated αS expression and purification

Codon-optimized αS in the pT7-7 vector was expressed in *E. coli* BL21 DE3. For acetylated αS, the N-terminal acetylation enzyme NatB from *Schizosaccharomyces pombe* was coexpressed in a second vector, pNatB (Johnson, et al., 2010). PNatB (pACYCduet-naa20-naa25) was a gift from Dan Mulvihill (Addgene plasmid # 53613). Expression was conducted in 50 mM phosphate-buffered 2YT-medium (pH 7.2) with 0.4% glycerol and 2 mM MgCl_2_, protein production was induced at OD 1-1.2 with 1 mM IPTG and run for 4 h at 37 °C. For ^15^N-labelled protein, αS or acetylated αS was expressed in M9 medium with 0.2% ^15^NH_4_Cl.

Sparsely labelled αS for DNP experiments was non-acetylated, expression was done in a similar way, in M9 medium using 0.4% [2-^13^C]-glucose and 0.2% ^15^NH_4_Cl. Isotope labelling of Phe, Gln, Glu, Pro, Asn, Asp, Met, Thr, Lys, and Ile was suppressed by supplementing sufficient quantities (150 μg/ml of each) of these unlabeled amino acids in the expression media as reported previously(Hong and Jakes, 1999).

Purification of αS or acetylated αS was carried out as previously described(Hoyer, et al., 2002), some changes to the original protocol have been made. Except for sparse labelled αS for which previous lysis in 20 mM Tris-HCl pH 8.0, 1 mM EDTA was done, cell lysis and release of αS or acetylated αS was performed by directly boiling the frozen cell pellet at 95 °C in a threefold volume of 20 mM sodium phosphate buffer, pH 7.4, for 30 min. Thermostable αS or acetylated αS remained in the supernatant after 30 min of centrifugation at 15000·*g* and 4 °C and was subsequently subjected to an ammonium sulfate precipitation by slowly adding saturated ammonium sulfate solution to 50% saturation at 4 °C. Precipitated protein was pelleted at 15000·g and 4 °C, dissolved in 50 mL of 50 mM Tris-HCl pH 8, sterile-filtered and loaded onto an ion exchange chromatography column (HiTrap Q FF, GE Healthcare), where αS or acetylated αS eluted at around 300 mM NaCl in 50 mM Tris-HCl pH 8. Elution fractions containing αS or acetylated αS were subjected to another ammonium sulfate precipitation and finally purified by a SEC run (Superdex 75 10/300, GE Healthcare) in 20 mM sodium phosphate pH 7.4, 50 mM NaCl.

N-terminal acetylation of acetyl-αS was checked by HPLC, mass spectrometry, and NMR, proved to be about 95% when coexpressed with NatB.

### Membrane scaffold proteins expression and purification

As reported in before(Bayburt, et al., 1998) *E. coli* BL21 (DE3) were transformed with MSP1D1 or MSP1D1ΔH5 plasmid DNA in vector pET28a. Cells were grown in LB medium, induced by 1 mM IPTG at an optical density of 0.7 and incubated 5-6 hours at 37°C, then pelleted down. Cells were resuspended in 50 mM Tris-HCl pH 8, 500 mM NaCl (buffer B) supplemented with 6 M GdnHCl and EDTA-free Complete protease inhibitors (Macherey-Nagel) lysed by sonication (Bandelin Sonopuls MS72 probe), centrifuged at 17000·g for 1 hour (Beckman J2-21 rotor JA-20.1) and incubated 1 hour with previously equilibrated 2.5 ml Ni-NTA agarose resin/3L culture (Macherey-Nagel). Column was washed with 4 CV buffer B, 4 CV buffer B supplemented with 1% Triton X-100, 4 CV buffer B + 60 mM Na-cholate, 4 CV buffer B, 4 CV buffer B + 20 mM imidazole. Four fractions of 1 CV were eluted with 250 mM imidazole. The whole process was kept at 4°C in a cold room. The elution fractions were pooled and dialysed against 100-fold 200 mM Tris-HCl pH 7.5, 100 mM NaCl. N-terminal His-tag was cleaved using TEV protease incubated overnight at 4 °C. ΔHis-MSP was separated from MSP by IMAC and concentrated to the desired molarity using a Vivaspin centrifugal device of 10 kDa MWCO.

### Nanodiscs assembly

Nanodiscs were assembled according to established protocols(Bayburt, et al., 2002). In short, lipids chloroform stocks were dried under nitrogen flow to obtain a lipid film and stored under vacuum overnight. ΔHis-MSP1D1 or MSP1D1ΔH5 and the appropriate amount of lipids (Avanti Polar Lipids) solubilized in 60 mM Na-cholate were mixed together in 20 mM Tris-HCl pH 7.5, 100 mM NaCl, 5 mM EDTA. The scaffold-to-lipids molar ratio was calculated from geometrical considerations. 20% w/v of previously washed Biobeads SM-2 (Biorad) were added and the mixture incubated at room temperature overnight. The Biobeads were removed by centrifugation and once again 20% w/v were added for an additional 4-5 hours. Finally, they were purified by SEC on a HiLoad 16/600 Superdex 200 pg column (GE Healthcare) equilibrated with 20 mM sodium phosphate pH 7.4, 50 mM NaCl using a Äkta pure device at a flow rate of 1 ml/min. The quality of NDs preparation was check by the SEC chromatogram as well as by DLS (PSS Nicomp). NDs were concentrated to the desired molarity using a Vivaspin centrifugal device of 10 kDa MWCO.

### Bio-layer interferometry (BLI)

NDs were immobilized on the sensor surface of amine reactive biosensors (AG2R) (fortéBIO, PALL Life Science) after EDC/NHS activation to a final level between 1.2 and 1.8 nm depending on the NDs type using an Octet RED96 instrument (fortéBIO, PALL Life Science). All biosensors were quenched with 1 M ethanolamine for 3 min. All experiments were carried out in multi cycle kinetics at 25 °C. Association of αS in running buffer (20 mM sodium phosphate pH 7.4, 50 mM NaCl) on NDs and reference biosensors was recorded for 120 s, followed by a dissociation phase of 360 s. Sensorgrams were double referenced using the reference biosensors and a buffer cycle. Steady-state analysis was realized by fitting the □ αS concentration dependency of the highest response against with a simple 1:1 binding model. After normalization, all on and off curves were fitted against simple exponential build-up or decays and led to similar on- and off-rates.

### Solution NMR spectroscopy

Solution NMR experiments were performed on a Bruker Avance III HD spectrometer operating at 600 MHz ^1^H Larmor frequency, equipped with a triple resonance TCI (^1^H, ^13^C, ^15^N) cryoprobe and shielded z-gradients. If not stated otherwise, all experiments data were recorded at 10 °C with an αS concentration of 50 μM in 20 mM sodium phosphate pH 7.4, 50 mM NaCl, 10% (v/v) ^2^H_2_O and ND concentration was set to 25 μM (one αS per membrane leaflet). All [^1^H-^15^N]-TROSY-HSQC NMR spectra were acquired with 32 scans and 256 indirect increments, processed with TOPSPIN 3.2 (Bruker) and analyzed with CCPN(Vranken, et al., 2005). Peaks were automatically integrated and the ratio of volumes in the presence and absence of NDs plotted against the primary sequence. Outliers as results of peak overlap and/or ambiguities were removed.

### ThT fluorescence aggregation assays

50 μM of αS or acetylated αS was mixed with either 25 μM (2:1), 3.125 μM (16:1) or 0.781 μM (64:1) nanodiscs with different lipid mixtures. Assays were conducted in 20 mM sodium phosphate buffer pH 7.4 or 20 mM acetate buffer pH 5.3 with 50 mM NaCl, 0.02% NaN_3_ and 10 μM Thioflavin T. Unless otherwise stated, triplicates of 120 μl were pipetted into 96-well half area well plates with non-binding surface (Corning No. 3881, black, clear bottom) containing a glass ball for mixing and incubated at 37 °C for up to 7 days. Orbital shaking at 217 rpm was used for 15 s every 20 min. Thioflavin T fluorescence was excited at 445 nm and measured at 485 nm every 20 min with 15 s of shaking prior to the measurement in a plate reader (Tecan Spark 10M or Tecan infinite M1000PRO).

For the seeded experiments, fibril seeds of αS or acetylated αS were prepared as follows: 300 μl of 100 μM αS or acetylated αS was fibrillated at 37 °C and 800 rpm for 3 days in a 2 ml tube containing a glass ball in a Thermomixer (Eppendorf). The fibril solution was diluted to 50 μM and sonicated with a tip sonicator (Bandelin Sonopuls HD3200, BANDELIN electronic) at 10% power (20 W) for 60 s, with 1 s pulses on and 4 s off in between. Seed solution was diluted 20-fold for the aggregation assays (2.5 μM, 5%).

Kinetic curves were corrected by subtracting the curve of buffer (containing NDs) in the presence of ThT and normalized to highest fluorescence intensity (in line with comparable fibril mass seen in SDS-PAGE after the aggregation assay). The corresponding triplicates are shown as transparent circles in order to visualize the reproducibility of each experiment. In the case of quiescent nucleation and seeded assays no normalization was applied and data were recorded without the presence of glass balls and without plate shaking.

### Sodium Dodecylsulfate – Polyacrylamide Gel Electrophoresis (SDS-PAGE)

In order to compare the amounts of soluble and fibrillated αS or acetylated αS in the aggregation samples, 100 μl of each triplicate sample were taken out of the well plate, combined in 1.5 ml tubes and spun down at 20000·g and 20 °C for 30 min. Supernatants (~290 μl) was removed and pellets were resuspended in 280 μl buffer and SDS-sample buffer (4-fold) was added. Samples were boiled for 15 min at 98 °C and subsequently 10 μl were loaded onto a 15% SDS-gel together with standards of αS or acetylated αS and nanodiscs.

### Dynamic Nuclear Polarization (DNP) NMR spectroscopy

Magic-angle spinning solid-state DNP experiments were performed on a Bruker Avance III HD spectrometer operating at 600 MHz, equipped with a 395.18 GHz second-harmonic gyrotron and a 3.2 mm ^1^H, ^13^C, ^15^N triple resonance low-temperature MAS probe. Data were collected at 100 K, 9 kHz MAS speed and 9 W continuous-wave microwave power. The samples were prepared from sparsely labelled non-acetylated αS (250 μg) in the presence or in the absence of 2:1 molar ratio of 100% POPG nanodiscs and filled into 3.2 mm sapphire rotors. Final sample conditions were 15 mM sodium phosphate pH 7.4, 25 mM sodium chloride, 30% ^2^H_2_O, 60% glycerol-d_6_ and 2.5 mM AMUPOL(Sauvee, et al., 2013). Two-dimensional [^13^C-^13^C]-Proton-Driven Spin Diffusion (PDSD) experiments with 1 s mixing time were performed. ^1^H decoupling using SPINAL64 with a decoupling field of 104 kHz was employed during evolution and detection periods. Both experiments were conducted using 300 *t*_1_ increments with 16 and 48 scans each for αS in the absence and in the presence of nanodiscs, respectively. A recycle delay of 5 s was used in both experiments. Both spectra were processed using Topspin 3.2 (Bruker) using identical parameters with squared and shifted sine bell function (qsine 2.5) for apodization.

### Molecular Dynamics simulations

As starting conformation for the MD simulations, the NMR structure of micelle-bound αS (PDB 1XQ8) was used, considering only the first 61 residues in order to concentrate on the membrane binding region of αS. The Amber99sb-ILDN force field (Lindorff-Larsen, et al., 2010) was used for αS, which was simulated in its non-acetylated form (i.e., with NH3+ at the N-terminus) and with a C-terminal N-methyl amide capping group to account for the fact that αS would continue beyond residue 61. All lysine side chains were modeled as positively charged, glutamate and aspartate as negatively charged, while glutamine and histidine residues were considered to be neutral corresponding to a pH of 7.4. The protein was placed either 0.5 nm or 1.5 nm above the membrane surface. A starting orientation with the negatively charged side chains pointing away from the membrane and the lysine side chains being oriented towards the membrane surface were chosen (Fig. 3d). For modeling the lipid bilayer, membrane patches consisting of POPC/POPG (1:1) or DMPC/DMPG (1:1) involving 512 lipids (256 lipids per leaflet) were built using CHARMM-GUI (Lee, et al., 2016) and modeled with Slipids force field parameters (Jambeck and Lyubartsev, 2012, Jambeck and Lyubartsev, 2013). Before αS was added, both lipid bilayers were solvated and simulated for 500 ns (POPC/POPG) or 1000 ns (DMPC/DMPG) to obtain relaxed membranes. Here, the same simulation procedure was employed as described below. αS was placed above the membrane, the protein-membrane complex solvated using the TIP3 water model, Na+ and Cl-added to neutralize the system and to mimic the Na+ concentration used in the experiments. The ion parameters of Smith and Dang (Smith and Dang, 1994) were used. The system was then subjected to steepest descent energy minimization, followed by MD equilibration in the NVT ensemble for 1 ns at 10 °C using the V-rescale thermostat (Bussi, et al., 2007) with a time constant of 0.5 ps and separate temperature coupling for the protein, membrane and water/ions. Afterwards, 1 ns of NPT equilibration was performed using the Nose-Hoover thermostat (Hoover, 1985, Nosé, 1984) and Parrinello-Rahman barostat (Parrinello and Rahman, 1981) with semiisotropic pressure scaling, a reference pressure of 1 bar, a time constant of 10.0 ps and an isothermal compressibility of 4.5 × 10-5 bar-1. During both equilibration steps, restraints were applied to the positions of the P-atoms of the lipids and terminal C-atoms of their tails with a force constant of 1000 kJ mol-1 nm-2. All bond lengths were constrained using the Lincs algorithm (Hess, et al., 1997). The Coulombic interactions were calculated using the Particle mesh Ewald (PME) method (Darden, et al., 1993, Essmann, et al., 1995) with a cut-off of 1.0 nm for the short-range interactions and a Fourier spacing of 0.12 nm. The cut-off for the van der Waals interactions was set at 1.4 nm. Periodic boundary conditions were employed in all directions. For the MD production runs the same settings as for the NPT equilibration were used, except that all position restraints were removed. All MD simulations were performed at 10 °C with a time step of 2 fs for integration using the GROMACS 4.6 molecular dynamics package (Hess, et al., 2008). For the analysis, which was performed using Gromacs and Membrainy tools (Carr and MacPhee, 2015), only the last 250 ns of each production run was used. An overview of the production runs, can be found in Figure 3 – figure supplement 2.

### Differential Scanning Calorimetry (DSC)

Samples of approximately 5 μM NDs of different types (and if stated 10 μM αS) were degassed for at least 20 min at 30 °C and measured in a Microcal VP-DSC instrument (Malvern Instruments). The thermograms were acquired up-scan from 5°C to 45°C at a scanning rate of 0.5°C/min and corrected by subtracting the thermogram of buffer.

### Atomic Force Microscopy (AFM)

AFM images were taken in air, using a Nanowizard III atomic force microscope (JPK, Berlin). Samples were taken at the end of aggregation experiments and diluted in pure water to ca. 1 μM protein concentration. 10 μl were added onto freshly cleaved mica and left to dry. Then, they were gently rinsed with water to remove excess salt. Imaging was performed using tapping mode with NSC 36 cantilevers (MikroMasch), with resonant frequencies between 70 and 150 kHz.

## ACKNOWLEDGMENTS

The authors acknowledge access to the Jülich-Düsseldorf Biomolecular NMR Center and thank the CSC – IT Center for Science, Finland, for computational resources. Financial support from an Emmy Noether grant of the DFG ET103/2-1 to M.E. and an ERC consolidator grant to W.H. is gratefully acknowledged. T.V. and M.M.W. acknowledge support from the International Graduate School of Protein Science and Technology (iGRASP_seed_) granted by the Ministry of Innovation, Science and Research of the state North Rhine-Westphalia. This work was supported by the DFG (HE 3243/4-1) and by the Ministry of Innovation, Science and Research within the framework of the NRW Strategieprojekt BioSC (BioSc seed fund). We thank Dr. Aldino Viegas for help with NMR data acquisition and Dr. Céline Galvagnion for fruitful discussions.

## COMPETING INTERESTS

The authors declare that no competing interests exist.

## AUTHOR CONTRIBUTION

H.H., B.S., D.W., W.H., A.K.B. and M.E. designed the experiments. T.V., M.M.W., H.S., B.U. and C.P. performed the experiments. T.V. and M.M.W. analyzed the data. T.V. and M.E. wrote the manuscript. All authors commented on the manuscript.

## FIGURE SUPPLEMENT

**Figure 1 – figure supplement 1.**
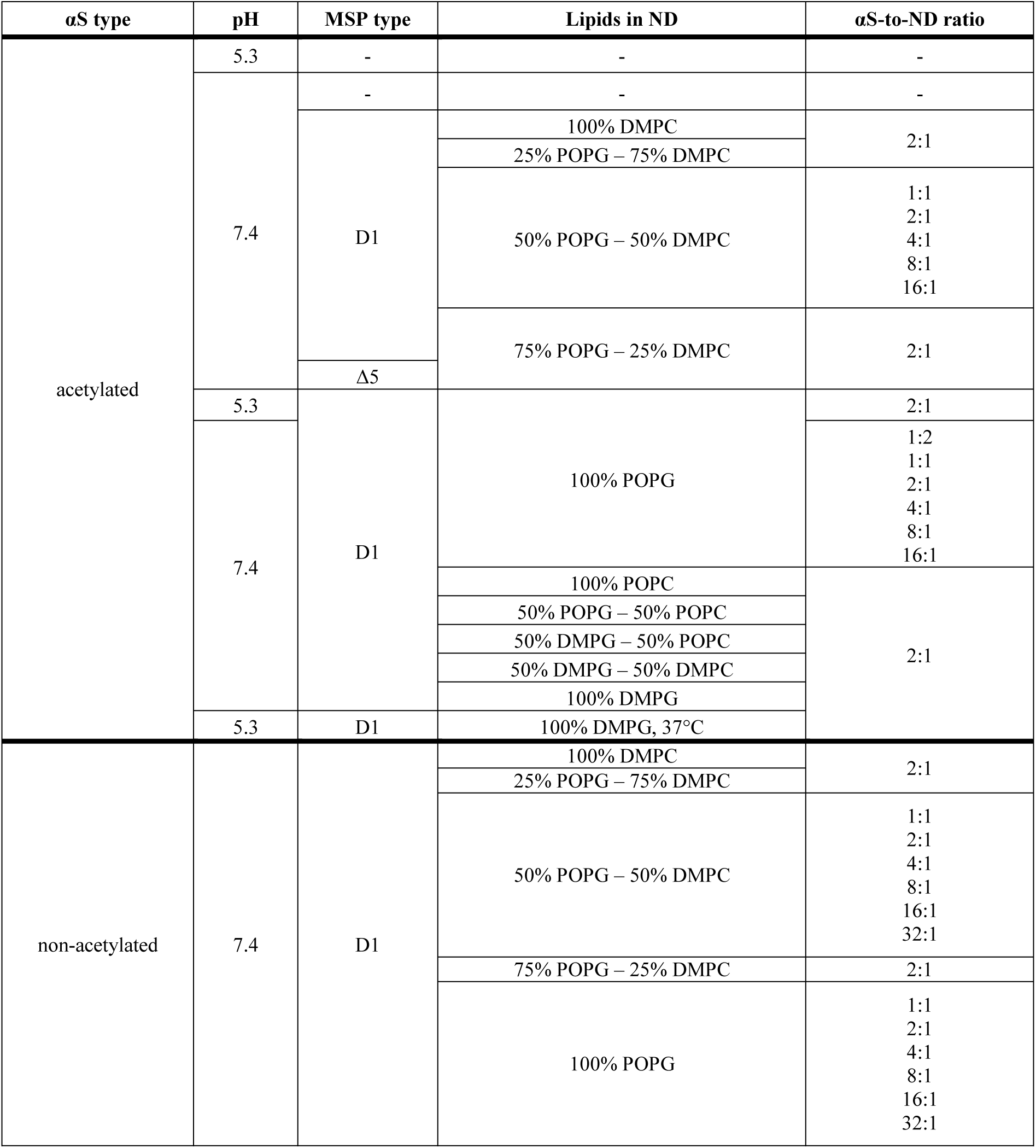
Summary of NMR samples used in the study.

**Figure 1 – figure supplement 2.**
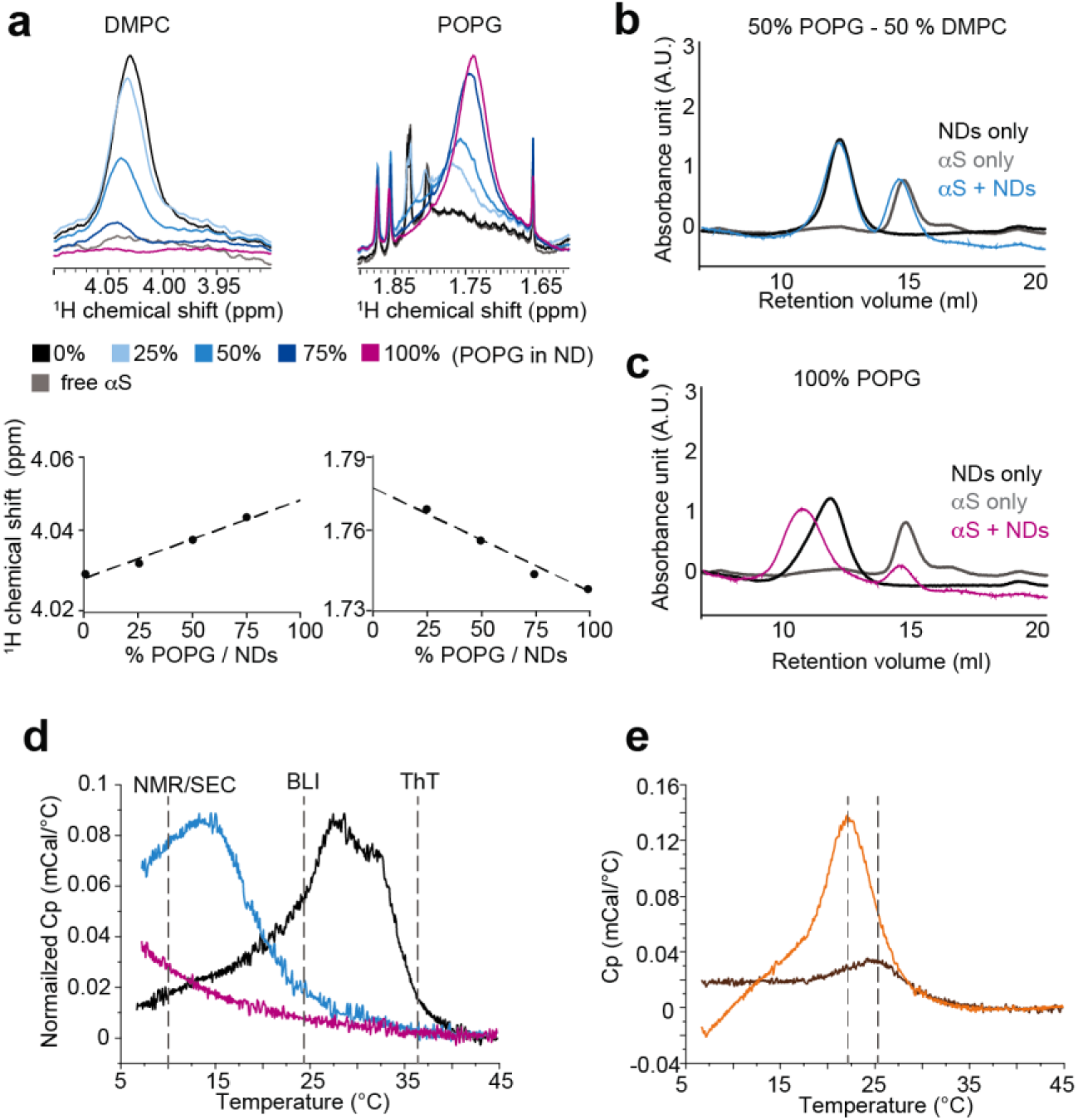
Formation of lipid mixtures in NDs, lipid phase transitions and stability of αS association. (**a**) NMR ^1^H-1D spectra of αS in the presence of NDs containing different POPG-to-DMPC ratios corresponding to the samples shown in Fig. 1a-b. Chemical shifts ranges for signals from either DMPC (left, choline methyl groups) or POPG (right, ^1^H next to the unsaturation) are displayed. The volumes of the peaks reflect within an error of approximately 10% the aimed lipid compositions. In addition to volume changes also chemical shifts perturbations for both lipid specific peaks are visible and follow a rather linear dependence on the composition. The NMR data therefore report on both the presence and mixing of both lipid types in single nanodiscs. (**b**) Size exclusion chromatography (SEC) profiles (Superdex 200 13/300 gl, GE Healthcare) of αS alone (grey), 50% POPG NDs alone (black) and both after mixing (same amounts as in the isolated case) and incubation for 24h at room temperature (blue). The clear separation of αS from NDs in the mixture points to a fast-kinetic exchange between free and bound αS. The conserved total absorbance additionally confirms the stability of the NDs in the presence of αS. (**c**) Same as in (b) but using 100% POPG NDs. The SEC profile of the mixture of αS with 100% POPG NDs shows a clear reduction of free αS and a size increase of the ND peak, pointing to a strong interaction between the two components. (**d**) Differential scanning calorimetry (DSC) thermograms of nanodiscs prepared with either 100% DMPC (black), 100% POPG (purple) or a 50%-50% mixture thereof (blue), showing phase transitions temperatures around 28°C, below 5°C and 13°C, respectively. Temperatures at which different measurements were performed are highlighted. (**e**) DSC profile of 100% DMPG NDs in the absence (brown) or presence of αS (orange). While the presence of αS leads to a lower phase transition temperatures of the lipid bilayer, in line with what was reported using SUVs, the transition temperature differences is much smaller for NDs (ΔTm=2.5 °C) as compared to SUVs (ΔTm=11 °C) in line with a largely increase stability of the NDs.

**Figure 3 – figure supplement 1.**
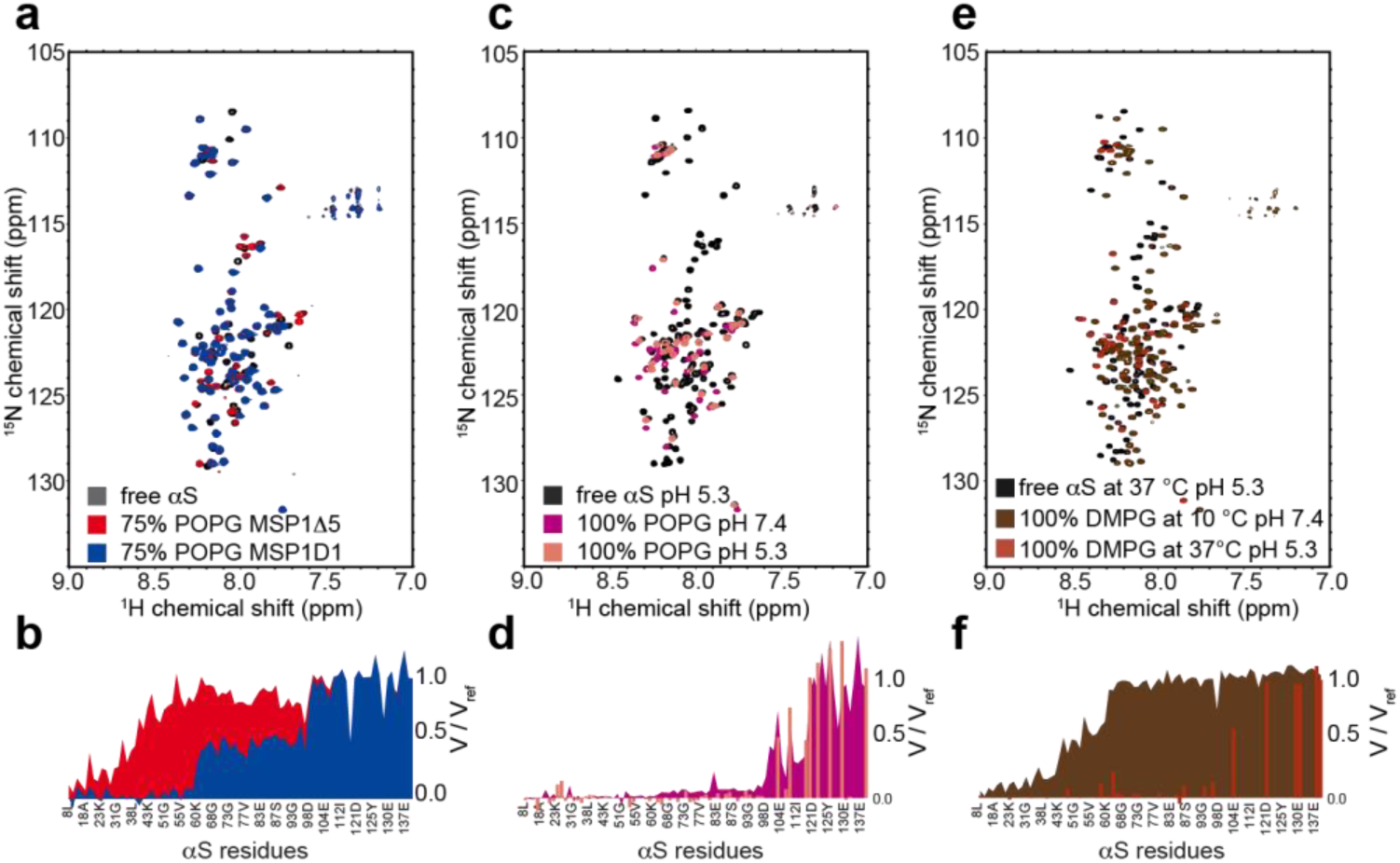
αS-ND interactions using different MSP constructs, lower pH and higher temperature. NMR spectra (**a**) and the respective attenuation profiles (**b**) of [^15^N]-acetylated-αS (50 μM) in the absence or in the presence of 25 μM NDs with 75% POPG content, assembled using the regular MSP1D1 construct (blue) or with the smaller MSP1Δ5 (red). Spectra (**c**) and attenuation profiles (**d**) corresponding to the binding of αS to NDs containing 100% POPG lipids at pH 7.4 (purple) and pH 5.3 (pale orange). Transferable assignments (bars) show that no significant difference in binding mode is visible upon pH variation. Spectra (**e**) and attenuation profiles (**f**) corresponding to the binding of αS to NDs containing 100% DMPG lipids in their gel phase (10°C, brown) or their fluid phase (37°C, red). Since normally measurement at 37 °C leads to considerable peak loss of N-terminal αS residues due to water exchange processes, the spectrum was recorded at pH 5.3 (counteracting water exchange). Note that the pH shift alone has no significant effect on binding (see (**c**) and (**d**)). Reference spectra of respective free αS are shown in dark grey.

**Figure 3 – figure supplement 2.**
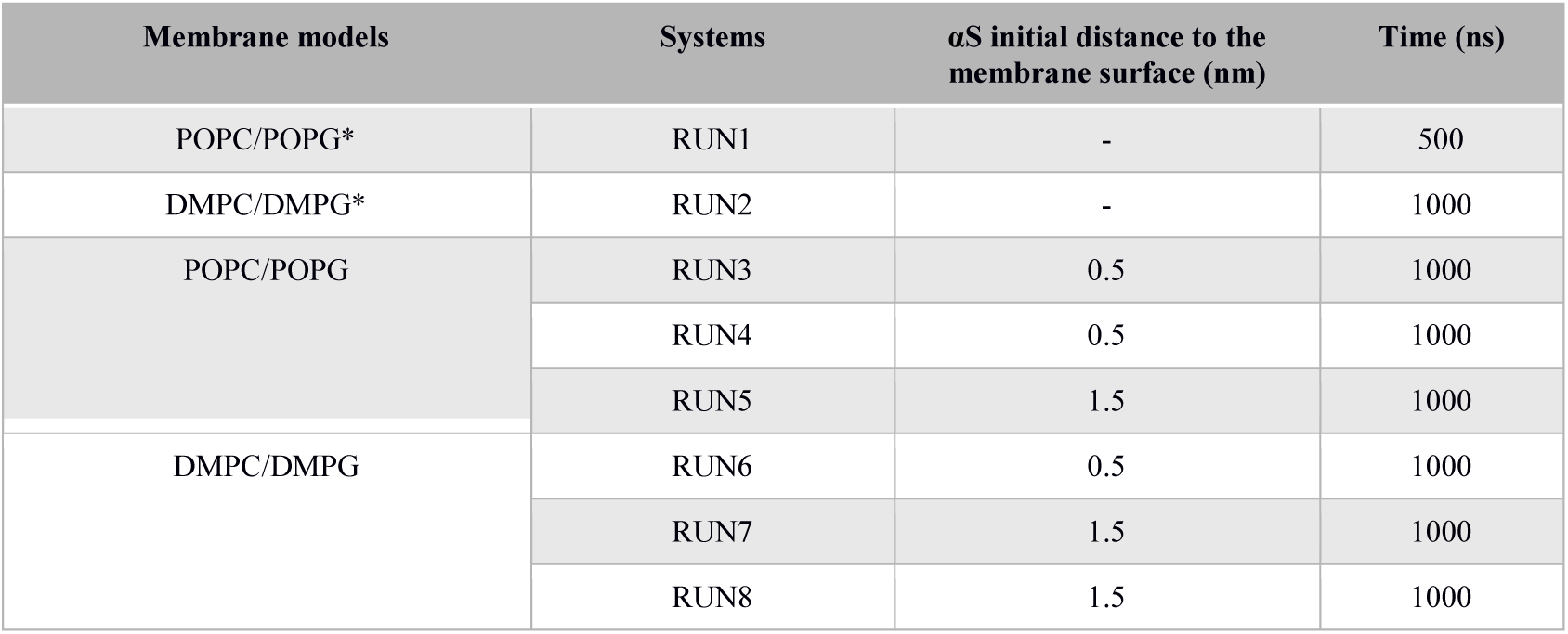
Summary of MD simulations used for the study.

**Figure 3 – figure supplement 3.**
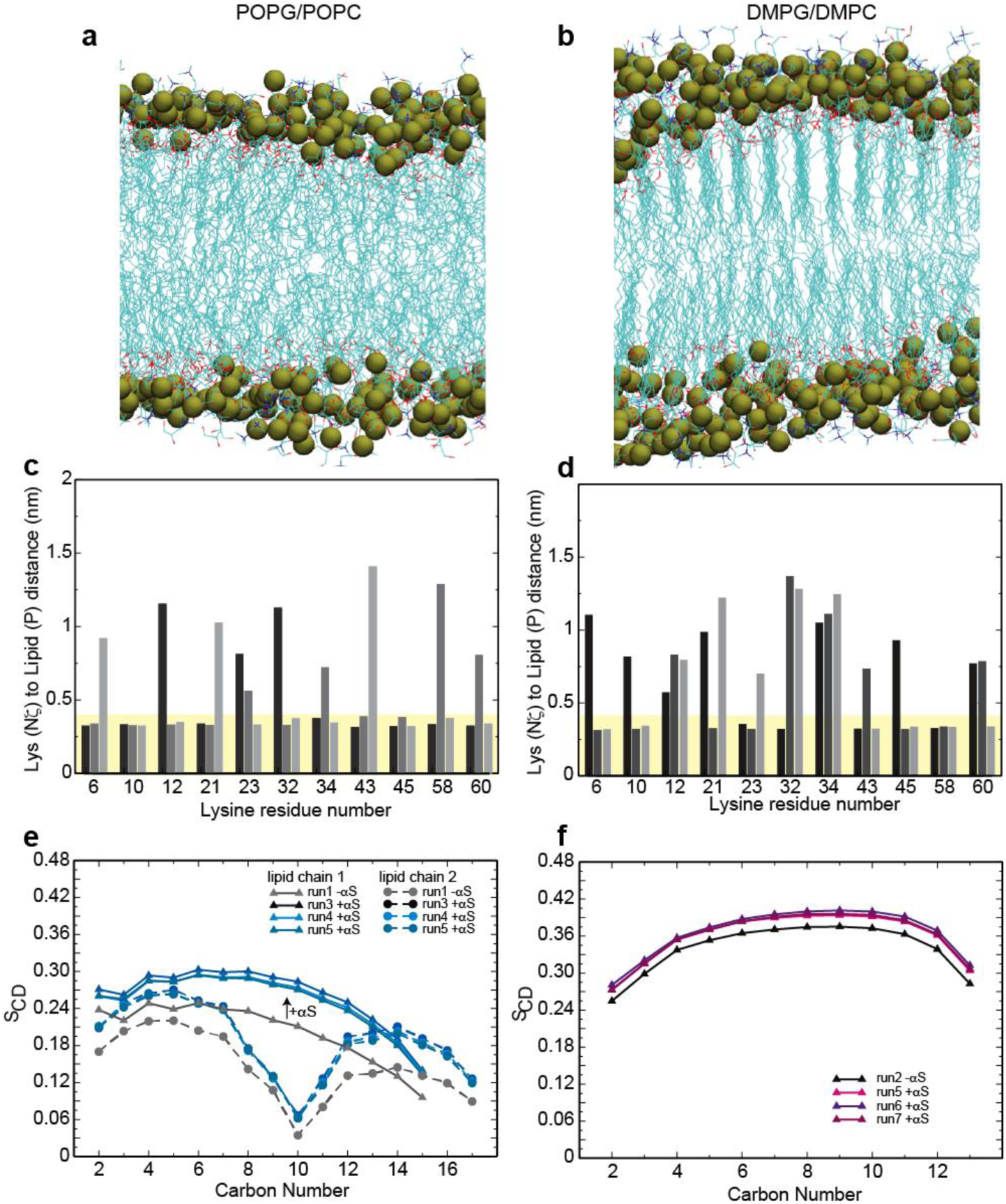
Lipid properties and αS-membrane interactions as seen by MD simulations. (**a,b**) Molecular arrangement of lipid bilayer in the absence of αS for POPG/POPC lipids in fluid phase (**a**) and DMPG/DMPC lipids in gel phase (**b**) (end of run1/run2, Table S2). (**c,d**) Distances for indicated lysine residues from Lys-Nξ to the phosphorus atom of the next anionic lipid as found in different runs of MD simulations for POPG/POPC (c, individual bars for run3-5, Table S2) and DMPG/DMPC (d, individual bars for run5-7, Table S2) membranes. Distances that could promote lipid-mediated salt bridges are highlighted in yellow. (**e,f**) Calculated order parameters for indicated carbon atoms of the lipid fatty acids with and without αS for POPG/POPC (**e**) and DMPG/DMPC (**f**) membranes.

**Figure 4 – figure supplement 1.**
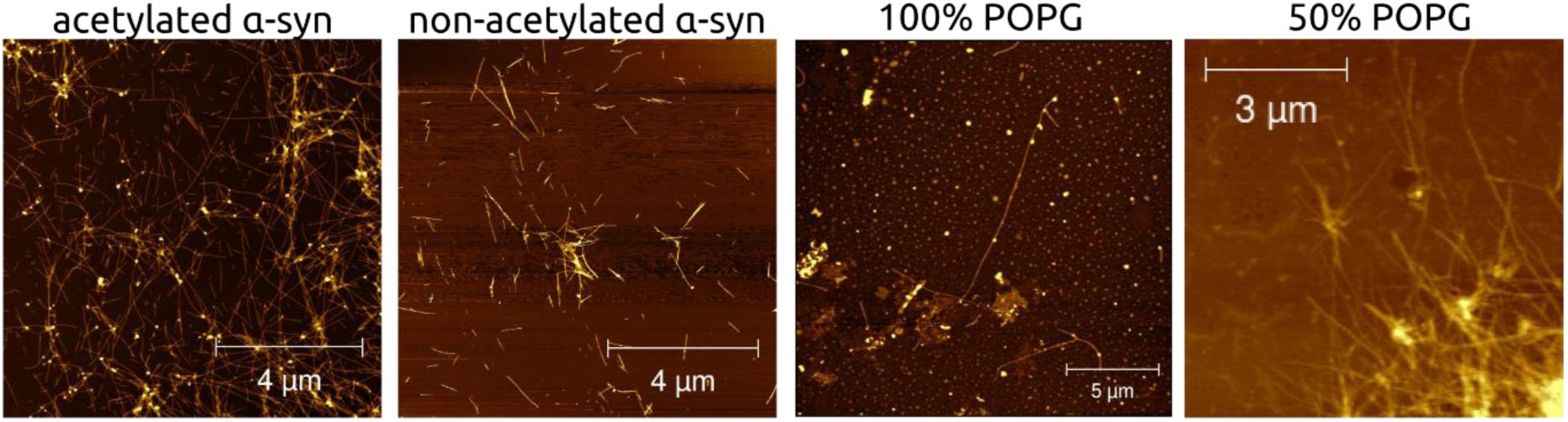
αS fibrils grown in the absence and presence of NDs have similar morphologies. AFM images of αS amyloid fibrils formed using acetylated αS and non-acetylated αS as well as acetylated αS in the presence of NDs with indicated lipid composition (from left to right).

**Figure 5 - figure supplement 1.**
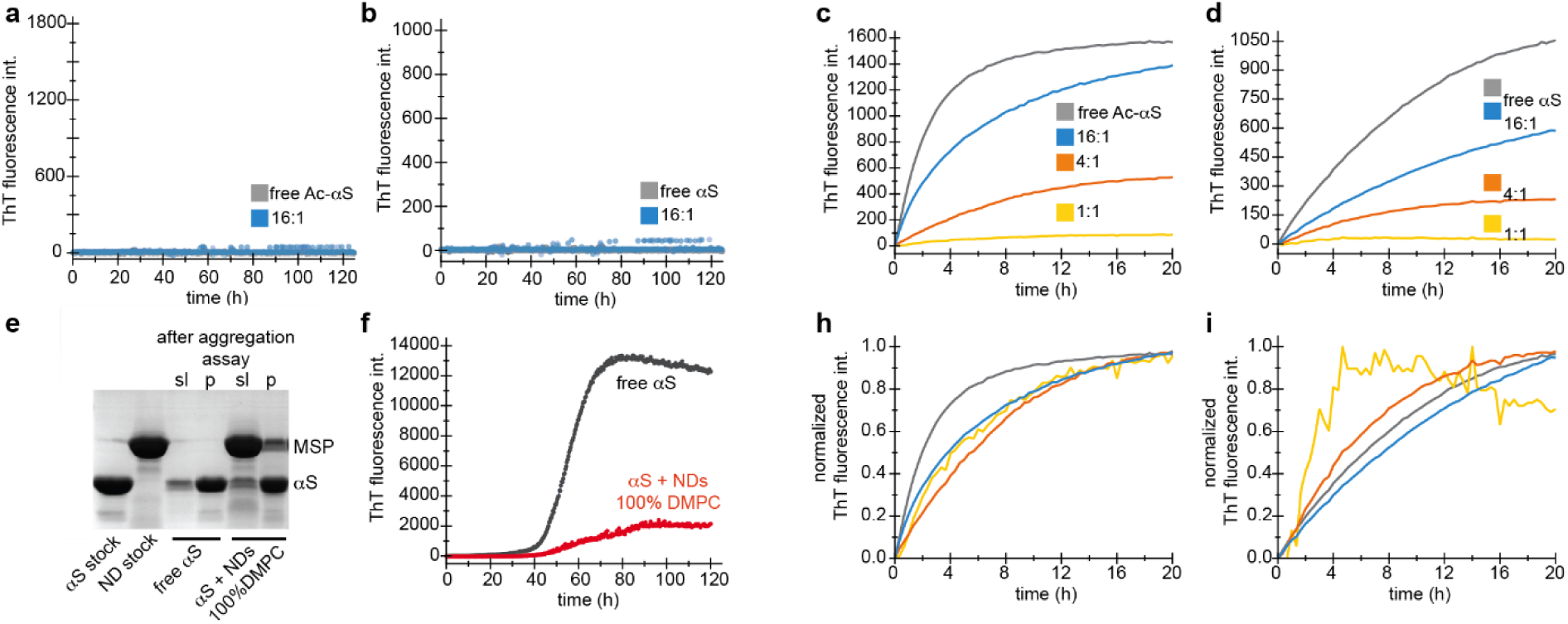
Amyloid fibril nucleation and elongation in the presence of 50% POPG ND and rational for normalization of ThT data. (**a,b**) Quiescent nucleation assay of acetylated αS (**a**) and non-acetylated (**b**) αS in the absence (grey) and presence of 50% POPG NDs (conditions analog to data shown in Fig. 4j,k, for 100% POPG NDs). (**c,d**,) Seeded aggregation assays under conditions analog to data shown in Fig. 4m,l, but using 50% POPG NDs. SDS-PAGE results (**e**) with ThT intensities (**f**) of identical samples. After the aggregation assay the αS-band for soluble (sl) and insoluble (p) protein show very similar intensities in the absence and presence of 100% DMPC NDs. Therefore, ThT intensities in the seeded aggregation assays were normalized (**h,i**).

